# Genome evolution and introgression in the New Zealand mud snails *Potamopyrgus estuarinus* and *Potamopyrgus kaitunuparaoa*

**DOI:** 10.1101/2023.10.31.565016

**Authors:** Peter D. Fields, Joseph R. Jalinsky, Laura Bankers, Kyle E. McElroy, Joel Sharbrough, Chelsea Higgins, Mary Morgan-Richards, Jeffrey L. Boore, Maurine Neiman, John M. Logsdon

## Abstract

We have sequenced, assembled, and analyzed the nuclear and mitochondrial genomes and transcriptomes of *Potamopyrgus estuarinus* and *Potamopyrgus kaitunuparaoa*, two prosobranch snail species native to New Zealand that together span the continuum from estuary to freshwater. These two species are the closest known relatives of the freshwater species *P. antipodarum—*a model for studying the evolution of sex, host-parasite coevolution, and biological invasiveness—and thus provide key evolutionary context for understanding its unusual biology. The *P. estuarinus* and *P. kaitunuparaoa* genomes are very similar in size and overall gene content. Comparative analyses of genome content indicate that these two species harbor a near-identical set of genes involved in meiosis and sperm functions, including seven genes with meiosis-specific functions. These results are consistent with obligate sexual reproduction in these two species and provide a framework for future analyses of *P. antipodarum—*a species comprising both obligately sexual and obligately asexual lineages, each separately derived from a sexual ancestor. Genome-wide multigene phylogenetic analyses indicate that *P. kaitunuparaoa* is likely the closest relative to *P. antipodarum.* We nevertheless show that there has been considerable introgression between *P. estuarinus* and *P. kaitunuparaoa.* That introgression does not extend to the mitochondrial genome, which appears to serve as a barrier to hybridization between *P. estuarinus* and *P. kaitunuparaoa.* Nuclear-encoded genes whose products function in joint mitochondrial-nuclear enzyme complexes exhibit similar patterns of non-introgression, indicating that incompatibilities between the mitochondrial and the nuclear genome may have prevented more extensive gene flow between these two species.

**Significance Statement:** No whole-nuclear genome sequences are currently available for snails of the genus *Potamopyrgus*, best known for *Potamopyrgus antipodarum*, an invasive species of rivers and lakes worldwide, and a famous model for the study of the evolution of sex. We have sequenced and analyzed the genome of sexual *P. estuarinus* and *P. kaitunuparaoa*, the closest known relatives of *P. antipodarum*. We show that 1) the genomes are very similar in gene content and size, 2) *P. kaitunuparaoa* is the closest relative to *P. antipodarum*, 3) significant introgression has occurred between *P. estuarinus* and *P. kaitunuparaoa*; these genomes set the stage for powerful direct analyses of the genomic features, *e.g*., sex to asexual transitions and invasive success, that make *P. antipodarum* unique.

## Introduction

Groups of closely related species or lineages that feature multiple separate transitions from one character state to another can be powerfully applied to answer general and trait-specific questions regarding evolutionary processes. Here, we present the first high-quality genome assemblies from two species of *Potamopyrgus*. This gastropod genus appears to represent a recent radiation across New Zealand (Haase 2008), and includes multiple examples of transitions from the ancestral ovoviparous and obligately sexual state to ovovivipary (Haase 2005) and obligately asexual lineages (Dybdahl and Lively 1995; Paczesniak, et al. 2013; Wallace 1992). These transitions are of broad interest because both of these traits—the modes of egg production (oviparous *vs.* ovoviviparous) and of reproduction (sexual *vs*. asexual)—are likely amongst the most important predictors of the likelihood of success in particular environments (Bell 1982; Clutton-Brock 1991; Maynard Smith 1978). The genetic basis of transitions to asexual reproduction has been characterized at least in part in a handful of taxa (reviewed in Neiman, et al. 2014; more recent examples include Yagound, et al. 2020; Ma, et al. 2021; Mau, et al. 2021), but this information remains outstanding for the vast majority of asexual taxa. Transitions between ovipary and viviparity (live birth, equivalent to ovoviviparity) have received quite a bit of attention in vertebrates, but comparatively little is known in invertebrates such as gastropods (Mamos, et al. 2021). One pattern that has emerged from studies of ovipary-vivipary transitions in molluscs is that derived viviparity is relatively common in freshwater taxa, while ancestral oviparity is the norm in marine species (Glaubrecht 2006; Köhler, et al. 2004).

Our focus here is to provide and analyze the first genome assemblies from two *Potamopyrgus* species: *Potamopyrgus estuarinus* and *P. kaitunuparaoa*. These are amongst the closest known evolutionary relatives of *P. antipodarum* (Haase 2008), itself an emerging model species for studying the evolution of asexual reproduction (*e.g.*, McElroy, et al. 2021): multiple (>>10) sexual to asexual transitions have occurred within *P. antipodarum* (Dybdahl and Lively 1995; Paczesniak, et al. 2013). Furthermore, a genome sequencing project is underway for *P. antipodarum,* including representatives from both sexual and asexual lineages (McElroy, et al. 2021; Sharbrough, et al. 2023; Sharbrough, et al. 2018). By completing the genomes of *P. estuarinus* and *P. kaitunuparaoa*, two closely related taxa with the apparently ancestral character states of obligately sexual reproduction involving the production of oviparous eggs, a clear framework is now laid for identifying the underpinnings of reproductive evolution in *Potamopyrgus.* Indeed, these genome assemblies provide key resources required for more detailed comparative analyses of *Potamopyrgus* to understand the genomic and genetic mechanisms involved in the repeated transitions to ovovivipary and asexuality in this genus. To this end, the careful inventory and detailed analyses of genes that are specifically involved in sexual reproduction are potentially key: biological processes such as meiosis and sperm production would be predicted to be under relaxed selective constraint (Schurko, et al. 2009). Thus, a clear picture of the ancestral states for such genes in a phylogenetically close and obligately sexual relative is both necessary and informative for identifying and understanding the genomic underpinnings—both cause and consequence—of sexual to asexual transitions.

In addition to our specific motivations to understand reproductive transitions, these new genomes contribute phylogenetic diversity to an otherwise sparse set of currently available mollusc genomes. Collectively, molluscs have a 500 million year fossil record, rich in biodiversity and replete with species of both scientific and economic importance. Despite their importance, genome resources for molluscan taxa, especially gastropods, have lagged compared to other bilaterian animals (Davison and Neiman 2021; Ghiselli, et al. 2021). As a result, functional genomics in molluscs has been forced to rely upon annotations ported over from model systems millions of years diverged from focal taxa. Phylogenomics can help span this divide by accounting for changes in rates and patterns of evolution across lineages that are often the hallmarks of altered function. For example, genes with relatively conserved functions are expected to exhibit relatively slow rates of evolution. By contrast, genes that gain new functions or that lose their ancestral functions are expected to exhibit relatively rapid rates of evolution. Thus, by identifying genes with relatively slow and relatively rapid rates of evolution, we can more accurately predict the putative functions of genes identified strictly by bioinformatic inference. We implemented such a comparative approach, using the transcriptomes of the close relative *P. antipodarum* and a more distantly related Litterinimorph snail, *Oncomelania hupensis*, as well as the genomes from two apple snails, *Pomacea canaliculata* and *Marisa cornuarietis*, to inform our genome annotations.

As a result of our detailed comparative analyses of these gastropod genomes, initially aimed at providing an accurate and informative genome assembly and annotation, we identified a substantial phylogenetic discordance among the three *Potamopyrgus* species between mitochondrial and nuclear gene trees. Careful analysis of these data indicates that *P. kaitunuparaoa* and *P. antipodarum* are sister taxa to the exclusion of *P. estuarinus* (matching the mitochondrial genome tree topology), but with substantial introgression between *P. estuarinus* and *P. kaitunuparaoa*. Notably, nuclear genes with predicted interactions with mitochondrially encoded genes were particularly resistant to introgression, indicating that mito-nuclear coevolution can contribute to barriers between species in *Potamopyrgus*.

In sum, we have assembled, annotated, and analyzed the genome sequences of two closely related snail species, *P. estuarinus* and *P. kaitunuparaoa.* These genomes add diversity to an otherwise phylogenetically depauperate collection of complete mollusc genomes, aiding future efforts to annotate and characterize genes in this important animal phylum. The *P. estuarinus* and *P. kaitunuparaoa* genomes reveal a clear picture of the ancestral state for sexual reproduction in this genome as a framework for assessing and understanding the evolution of asexuality in their close relative, *P. antipodarum.* Finally, phylogenetic analyses of these genomes reveal a substantial history of introgression associated with the divergence between *P. estuarinus* and *P. kaitunuparaoa*.

## Results & Discussion

We have sequenced, assembled, and annotated the genomes of *P. estuarinus* and *P. kaitunuparaoa.* Our analyses of these genomes demonstrate considerable similarity in genome size, structure, and gene content, including a conserved set of genes involved in meiosis. Phylogenetic analyses of both mitochondrial and nuclear genomes indicate a strongly supported sister relationship between *P. antipodarum* and *P. kaitunuparaoa,* but also reveal a substantial amount of introgression of nuclear genes between *P. estuarinus* and *P. kaitunuparaoa*. We describe and interpret these major findings in detail below.

### Genome Assembly, Annotation, and Assessment

Analyses from unassembled short reads indicate that these *Potamopyrgus* species have similar genomic characteristics. We estimated the haploid genome size for both *P. estuarinus* and *P. kaitunuparaoa* at around 500 Mb, with heterozygosity of 3.73% for *P. estuarinus* and 3.63% for *P. kaitunuparaoa* (Figure S2). Our assembled genome of *P. estuarinus* has a total length of 523.89 Mb (N=12,267 total contigs), an N50 of 80 Kb (N=1508), and a max contig length of 1.1 Mb. Because our *P. kaitunuparaoa* assembly is based on only short-read small-insert Illumina data, we could not obtain as contiguous of an assembly as for *P. estuarinus*. The inclusion of comparative scaffolding with our *P. estuarinus* resulted in the following assembly summary for *P. kaitunuparaoa*: total length of 597.46 Mb (N=132,386 total scaffolds and contigs), with an N50 of 48 Kb (N=2456), and a max contig length of 1.04 Mb. Overall, our assembly and annotation pipeline produced 32,238 protein-coding gene models for *P. estuarinus* and 30,311 protein-coding gene models for *P. kaitunuparaoa*.

Following the creation of the genome assemblies and resulting annotations for *P. estuarinus* and *P. kaitunuparaoa*, we used BUSCO v.5.4.2 to reevaluate biological completeness in both genomes. We compared both genomes to the metazoa_odb10 BUSCO reference gene set. We found that 84.6% of the 954 metazoa BUSCO genes were assembled in single copy in *P. estuarinus* (Figure 2c, S3a) and that 84.1% of the genes were assembled in single copy in *P. kaitunuparaoa* (Figure 2c, S3b). We used the BlobToolKit pipeline to assess the *P. estuarinus* and *P. kaitunuparaoa* genome assemblies for completeness and contamination (Figure S4). Scaffolds that were majority composed of bacterial, viral, or fungal sequence were identified as contaminants. Following the removal of these contaminant scaffolds, the assemblies were reassessed using BUSCO and BlobToolKit (Figures 2, S4).

**Figure 1.**
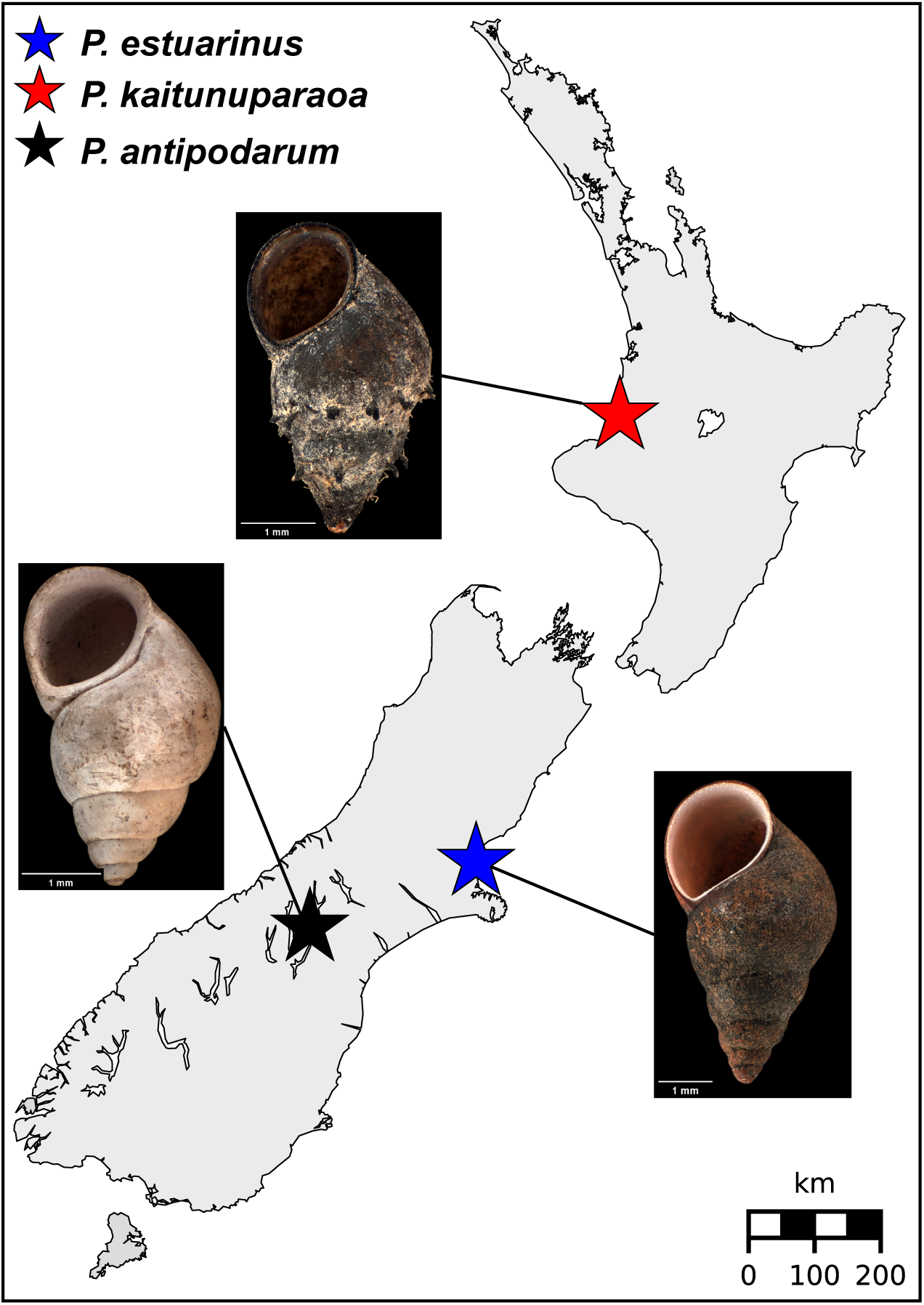
Map of New Zealand sampling locations of *P. estuarinus, P. kaitunuparaoa, and P. antipodarum*. Map locations of *P. estuarinus* (blue star), *P. kaitunuparaoa* (red star), and *P. antipodarum* (black star). Shell morphology for each species is depicted in associated images. Shell photographs were obtained from the Museum of New Zealand (Te Papa Tongarewa) and modified in accordance with the Creative Commons BY 4.0 license. Links to original photographs are provided in the Data Availability statement.

**Figure 2.**
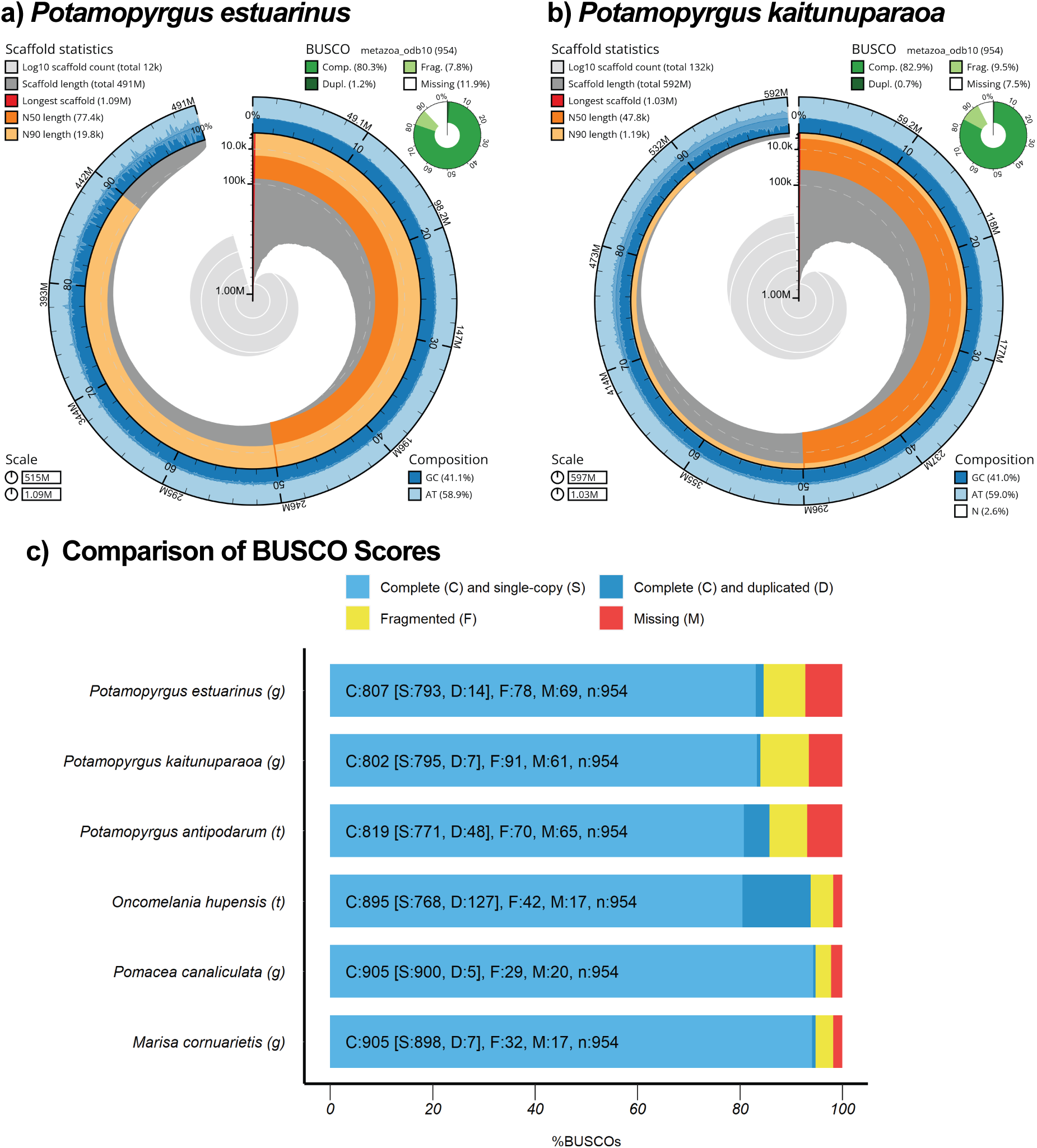
Genome assembly statistics for *P. estuarinus and P. kaitunuparaoa* genome assemblies. Snail plots describing genome assembly statistics for *P. estuarinus* (a) and *P. kaitunuparaoa* (a) after contaminant removal (*i.e*., sequences identifying with kingdoms in Bamfordvirae, Bacteria-undef, Viridiplantae or Fungi; or phylum in Pseudomonadota or Bacteroidota). The main plot in the center of each panel is divided into 1,000 size-ordered bins, with each bin representing 0.1% of the assembly (total length *P. estuarinus –* 514,633,712 bp; total length *P. kaitunuparaoa –* 596,959,456 bp). Sequence length distribution is shown in dark gray, with the radius scaled to the longest present sequence (red). The arcs in the plot represent the N50 sequence length (orange) and N90 sequence length (pale orange). The pale gray spiral shows the cumulative sequence count on a log scale, with orders of magnitude represented by white scale lines. GC, AT, and N percentages are reflected by the dark blue and pale blue area around the outside of the plot, in the same bins as the inner plot. A summary of BUSCO genes in the metazoa_odb10 dataset present after removal of contaminants are shown in the top right of each plot. BUSCO summary statistics for whole assemblies *P. estuarinus*, *P. kaitunuparaoa*, *P. antipodarum* (transcriptome), *O. hupensis* (transcriptome), *M. cornuarietis*, and *Po. canaliculata are shown in panel C*.

After contaminants were removed, BUSCO showed that a mean of 80.3% of metazoan markers were assembled in single copy in the *P. estuarinus* genome, with an updated N50 of 77.4 kb (Figure 2a). A total of 82.9% of metazoan BUSCO markers were assembled in single copy in the *P. kaitunuparaoa* genome. Following contaminant removal, *P. kaitunuparaoa* had an updated N50 of 47.8 kb (Figure 2b). Annotated transcripts from both genomes overwhelmingly showed significant homology, as assessed by BLAST similarity scores, to mollusc or other metazoan records (Figure S5a-d), particularly after potentially contaminating sequences were removed (Figure S5e-h). A detailed summary of genome statistics before and after the removal of contaminants can be found in Tables S1 and S2. Importantly, as can be seen with the BUSCO scores, our assemblies are likely highly biologically complete and thus informative for population and phylogenetic analysis. Our estimates of genomic repeat content using dnaPipeTE were also very similar across the two species, with 31.06% and 30.37% repeat portions of the *P. estuarinus* and *P. kaitunuparaoa* genome, respectively (Table S3). The estimated rDNA amounts for these two species fall within the range of rDNA content estimated from diploid sexual *P. antipodarum* using short reads to estimate repeat abundance (McElroy, et al. 2021).

### Comparative Genomic Assessment of Assembly and Annotation Quality

To evaluate the veracity of our gene models predicted from both *P. estuarinus* and *P. kaitunuparaoa* genomes, we inferred homologous proteins from independently assembled caenogastropod genomic and transcriptomic resources using OrthoFinder (Figure 3, Figure S6). We found 27,549 homologous gene groups that contained at least one protein from either *P. estuarinus* (representing all 32,238 *P. estuarinus* genes) or *P. kaitunuparaoa* (representing all 30,311 genes). The vast majority of homologous gene groups (103,672) were specific to either the *P. antipodarum* transcriptome or the *O. hupensis* transcriptome – these apparently spurious genes were not considered further, as they provided no comparative information. We found 1,713 homologous gene groups that contained exactly one protein from all six taxa (*i.e*.,there were 7,731 homologous gene groups for which all species were represented by at least one protein, representing 10,681 (33%) *P. estuarinus* genes and 10,438 (34%) *P. kaitunuparaoa* (see Figure 3). We also identified 6,614 gene groups that were single copy in *Potamopyrgus*, an additional 1,817 that were single copy in both *P. estuarinus* and *P. kaitunuparaoa* but multi-copy in the *P. antipodarum* transcriptome, and 12,202 homologous gene groups (representing 18,616 *P. estuarinus* genes and 19,097 *P. kaitunuparaoa* genes) that had at least one protein from all three *Potamopyrgus* assemblies. For *P. estuarinus*, we found that 26,001 (81%) of all *P. estuarinus* gene models were homologous to one or more proteins from at least one other independent assembly (*i.e*., excluding *P. kaitunuparaoa*). 25,150 (83%) *P. kaitunuparaoa* gene models satisfied the same criterion (*i.e*., excluding *P. estuarinus*). There were 1,513 homologous gene groups (representing 1,898 *P. estuarinus* genes and 1,940 *P. kaitunuparaoa* genes) that contained proteins from both *P. estuarinus* and *P. kaitunuparaoa* but no other assembly. We found 3,586 homologous gene groups (representing 4,338 genes) that were specific to *P. estuarinus* and 3,048 homologous gene groups (representing 3,220 genes) that were specific to *P. kaitunuparaoa*. These groups may represent false-positive gene annotations. There were no *P. estuarinus* or *P. kaitunuparaoa* proteins that were not assigned to homologous gene groups as a result of this process.

**Figure 3.**
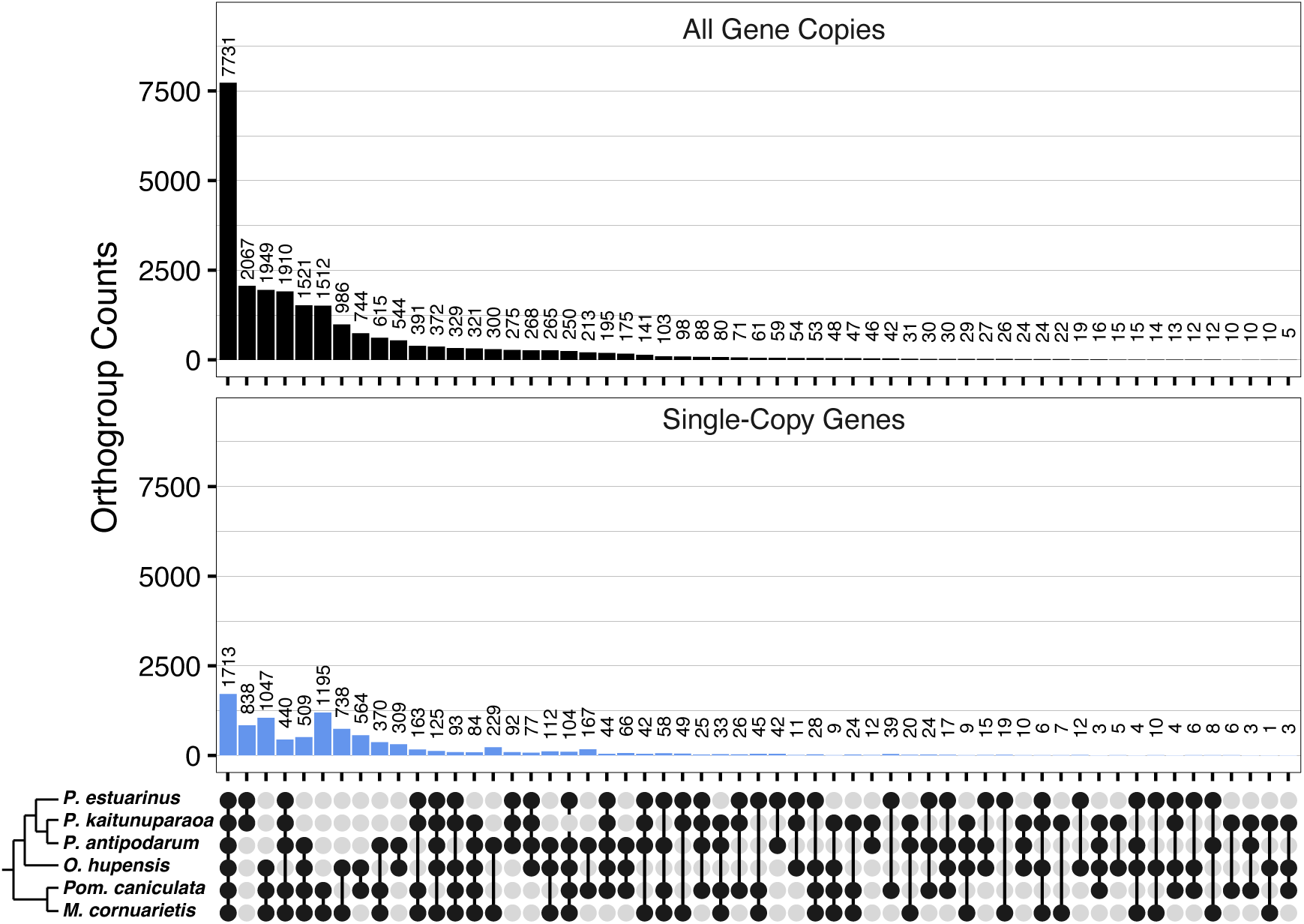
Homologous gene groups in assemblies from six caenogastropodan taxa. Upset plot depicting the number of genes fitting into homologous gene groups (group membership specified in bottom panel) for multi-copy genes (top panel) and single-copy genes (bottom panel). Singleton genes that are only present in a single assembly have been removed from this plot for display purposes.

To evaluate the quality of gene models, we estimated the rate of synonymous (*d_S_*) and nonsynonymous (*d_N_*) substitutions per site for 1,682 single-copy genes (excluding 30 genes that had internal stop codons in the *Pom. canaliculata* or the *M. cornuarietis* annotations) on all three possible tree topologies (*i.e*., *Pa-Pk*, *Pe-Pk*, and *Pa-Pe*), plus an analysis excluding *P. antipodarum* (to alleviate challenges caused by topological variation within *Potamopyrgus*), and an analysis including only the *Potamopyrgus* species (to alleviate challenges caused by saturation). Our gene models appear to be overwhelmingly consistent with being correctly annotated, as *d_N_* /*d_S_* estimates for the vast majority of genes (mean *d_N_* /*d_S_* +/- SD = 0.097 +/- 0.059; Figure S7) were low (*i.e*., << 1), as expected under a predominant force of purifying selection. The deep divergence between the apple snails and *Potamopyrgus* means that the observed overall synonymous substitution rates are highly saturated (mean *d_S_* = 4.35 synonymous substitutions per site). We can nevertheless use the branch-specific model of codeml (*i.e*., model 1) to estimate *d_S_* across the tree (Figure S8). Using this approach, and excluding 23 genes (1.4%) that were clear *d_S_* outliers (*i.e*., terminal branch *d_S_* values > 1), we found low *d_S_* rates (*i.e*., less than 10% divergence) within *Potamopyrgus* (mode *d_S_* +/- SD = 0.0783 +/- 0.10 synonymous substitutions per site), between *P. estuarinus* and *P. antipodarum* (mode *d_S_* +/- SD = 0.041 +/- 0.080 synonymous substitutions per site), and between *P. kaitunuparaoa* and *P. antipodarum* (mode *d_S_*= 0.041 +/- 0.083 synonymous substitutions per site) assuming the *Pa-Pk* tree. Similar estimates were obtained considering the other two tree topologies. In sum, analyses of molecular evolution indicate that *P. estuarinus* and *P. kaitunuparaoa* have high-quality gene model annotations and that divergence is extremely deep among *Caenogastropod* species that boast high-quality genomic resources.

### Analysis of Gene Families Associated with Meiotic Function

We inventoried and manually annotated the same 44 meiosis-related genes in three different molluscan gastropod species (*O. hupensis*, *M. cornuarietis*, and *Po. canaliculata*) in addition to the three focal *Potamopyrgus* species, to characterize the gain or loss of these genes in *Potamopyrgus* across a larger phylogenetic framework and to provide a curated assessment of our automated gene annotations. The presence of intact (complete and in-frame CDS) meiosis genes in both focal *Potamopyrgus* taxa indicates the capacity for sexual reproduction is maintained. This finding is of considerable importance in light of the coexistence of obligately sexual lineages with multiple separately derived obligately asexual lineages that characterizes *P. antipodarum*. Establishing a repertoire of genes involved in sexual reproduction (for sexual lineages) is a necessary prerequisite before focused comparisons across sexual and asexual lineages can take place (Schurko and Logsdon Jr 2008). The meiosis gene comparisons revealed several instances of taxa-specific gene loss and retention (Table 1). First, 17 of the 44 genes are present in the genome or transcriptome assemblies across all six species, and three genes (*CYCA, TIM2, RECQ3*) were absent from all six species. Three of the 44 genes were absent in all three *Potamopyrgus* species but present in all three other mollusks (*SEPARASE, RECQ2, and RECQ4*). *HOP1*, a meiosis-specific gene, is present only in *Po*. *canaliculata* and *M*. *cornuarietis*. *RAD51D*, a gene involved in homologous recombination and DNA repair, was found in all three *Potamopyrgus* taxa but was absent (or undetectable) from the other three species. That most, but not all, meiosis genes are present for each species and are not pseudogenes is not surprising in light of previous reports of taxon-specific meiosis gene loss in *bonafide* sexual species (Schurko and Logsdon Jr 2008; Malik, et al. 2008). In the current study, at least some of the apparent gene absences are likely a result of genome assemblies not correctly assembling the reads containing the genes, or in the case of the *O*. *hupensis* transcriptome, the absent genes were not being expressed in sufficient quantity to be captured by RNA sequencing. In short, the complement of meiosis genes in *Potamopyrgus* provides a framework for further investigation into how genes essential to sex evolve in an asexual background.

**Table 1.**
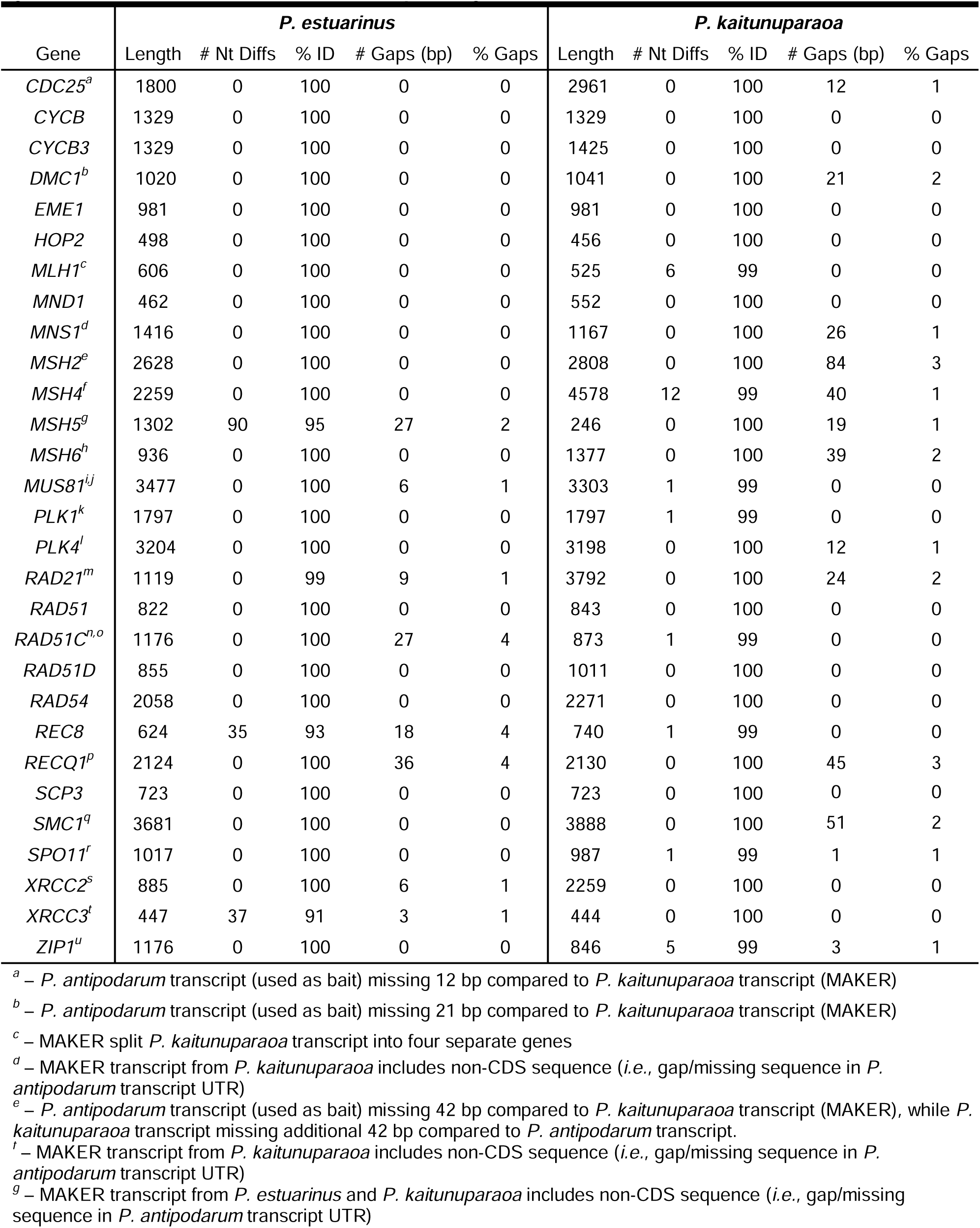
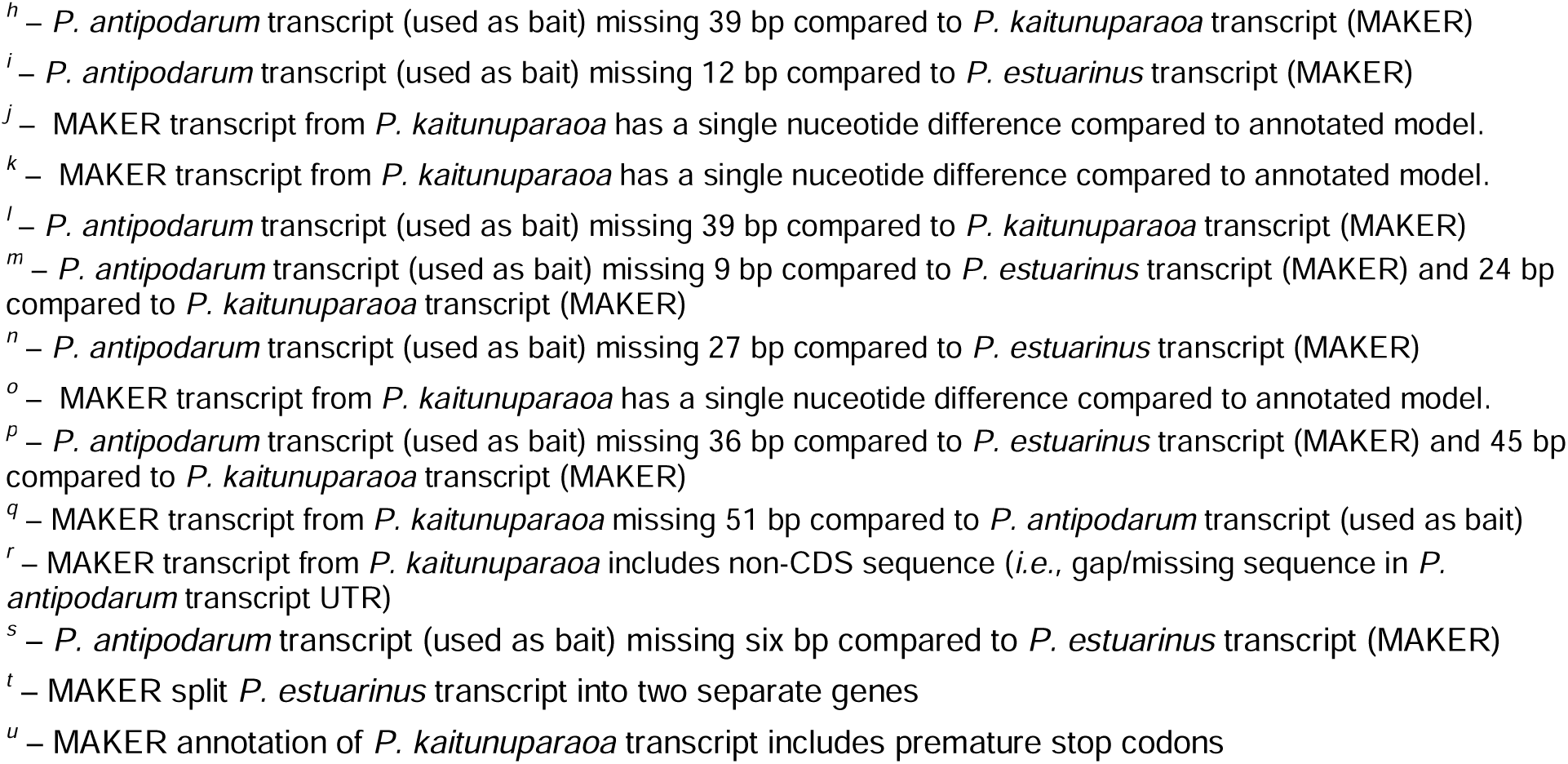
BLAST comparisons of the aligned portions of MAKER- and hand-annotated models of meiosis genes in the *P*. *estuarinus* and the *P*. *kaitunuparaoa* genome assemblies.

| Gene | *P. estuarinus* |  |  |  |  | *P. kaitunuparaoa* |  |  |  |  |
| --- | --- | --- | --- | --- | --- | --- | --- | --- | --- | --- |
|  | Length | # Nt Diffs | % ID | # Gaps (bp) | % Gaps | Length | # Nt Diffs | % ID | # Gaps (bp) | % Gaps |
| *CDC25<sup>a</sup>* | 1800 | 0 | 100 | 0 | 0 | 2961 | 0 | 100 | 12 | 1 |
| *CYCB* | 1329 | 0 | 100 | 0 | 0 | 1329 | 0 | 100 | 0 | 0 |
| *CYCB3* | 1329 | 0 | 100 | 0 | 0 | 1425 | 0 | 100 | 0 | 0 |
| *DMC1<sup>b</sup>* | 1020 | 0 | 100 | 0 | 0 | 1041 | 0 | 100 | 21 | 2 |
| *EME1* | 981 | 0 | 100 | 0 | 0 | 981 | 0 | 100 | 0 | 0 |
| *HOP2* | 498 | 0 | 100 | 0 | 0 | 456 | 0 | 100 | 0 | 0 |
| *MLH1<sup>c</sup>* | 606 | 0 | 100 | 0 | 0 | 525 | 6 | 99 | 0 | 0 |
| *MND1* | 462 | 0 | 100 | 0 | 0 | 552 | 0 | 100 | 0 | 0 |
| *MNS1<sup>d</sup>* | 1416 | 0 | 100 | 0 | 0 | 1167 | 0 | 100 | 26 | 1 |
| *MSH2<sup>e</sup>* | 2628 | 0 | 100 | 0 | 0 | 2808 | 0 | 100 | 84 | 3 |
| *MSH4<sup>f</sup>* | 2259 | 0 | 100 | 0 | 0 | 4578 | 12 | 99 | 40 | 1 |
| *MSH5<sup>g</sup>* | 1302 | 90 | 95 | 27 | 2 | 246 | 0 | 100 | 19 | 1 |
| *MSH6<sup>h</sup>* | 936 | 0 | 100 | 0 | 0 | 1377 | 0 | 100 | 39 | 2 |
| *MUS81<sup>i,j</sup>* | 3477 | 0 | 100 | 6 | 1 | 3303 | 1 | 99 | 0 | 0 |
| *PLK1<sup>k</sup>* | 1797 | 0 | 100 | 0 | 0 | 1797 | 1 | 99 | 0 | 0 |
| *PLK4<sup>l</sup>* | 3204 | 0 | 100 | 0 | 0 | 3198 | 0 | 100 | 12 | 1 |
| *RAD21<sup>m</sup>* | 1119 | 0 | 99 | 9 | 1 | 3792 | 0 | 100 | 24 | 2 |
| *RAD51* | 822 | 0 | 100 | 0 | 0 | 843 | 0 | 100 | 0 | 0 |
| *RAD51C<sup>n,o</sup>* | 1176 | 0 | 100 | 27 | 4 | 873 | 1 | 99 | 0 | 0 |
| *RAD51D* | 855 | 0 | 100 | 0 | 0 | 1011 | 0 | 100 | 0 | 0 |
| *RAD54* | 2058 | 0 | 100 | 0 | 0 | 2271 | 0 | 100 | 0 | 0 |
| *REC8* | 624 | 35 | 93 | 18 | 4 | 740 | 1 | 99 | 0 | 0 |
| *RECQ1<sup>p</sup>* | 2124 | 0 | 100 | 36 | 4 | 2130 | 0 | 100 | 45 | 3 |
| *SCP3* | 723 | 0 | 100 | 0 | 0 | 723 | 0 | 100 | 0 | 0 |
| *SMC1<sup>q</sup>* | 3681 | 0 | 100 | 0 | 0 | 3888 | 0 | 100 | 51 | 2 |
| *SPO11<sup>r</sup>* | 1017 | 0 | 100 | 0 | 0 | 987 | 1 | 99 | 1 | 1 |
| *XRCC2<sup>s</sup>* | 885 | 0 | 100 | 6 | 1 | 2259 | 0 | 100 | 0 | 0 |
| *XRCC3<sup>t</sup>* | 447 | 37 | 91 | 3 | 1 | 444 | 0 | 100 | 0 | 0 |
| *ZIP1<sup>u</sup>* | 1176 | 0 | 100 | 0 | 0 | 846 | 5 | 99 | 3 | 1 |

<sup>a</sup> – *P. antipodarum* transcript (used as bait) missing 12 bp compared to *P. kaitunuparaoa* transcript (MAKER)
<sup>b</sup> – *P. antipodarum* transcript (used as bait) missing 21 bp compared to *P. kaitunuparaoa* transcript (MAKER)
<sup>c</sup> – MAKER split *P. kaitunuparaoa* transcript into four separate genes
<sup>d</sup> – MAKER transcript from *P. kaitunuparaoa* includes non-CDS sequence (*i.e.*, gap/missing sequence in *P. antipodarum* transcript UTR)
<sup>e</sup> – *P. antipodarum* transcript (used as bait) missing 42 bp compared to *P. kaitunuparaoa* transcript (MAKER), while *P. kaitunuparaoa* transcript missing additional 42 bp compared to *P. antipodarum* transcript.
<sup>f</sup> – MAKER transcript from *P. kaitunuparaoa* includes non-CDS sequence (*i.e.*, gap/missing sequence in *P. antipodarum* transcript UTR)
<sup>g</sup> – MAKER transcript from *P. estuarinus* and *P. kaitunuparaoa* includes non-CDS sequence (*i.e.*, gap/missing sequence in *P. antipodarum* transcript UTR)

Another fundamental expectation for the loss of sex is that while genes critical for sex are under purifying selection and are evolving at similar rates in closely related sexual species, this constraint should be lifted for asexual lineages, resulting in a shift to relaxed selection for those lineages. Relaxed selection can act to degrade and render useless genes that are redundant in the absence of sex. That the majority of meiosis-specific genes remain intact in sexual *Potamopyrgus* species (Table 2) indicates that sex has been maintained and these genes are identifiable and retrievable. Indeed, for future studies of relaxed selection in the meiosis genes of asexual lineages, it is critical to establish that these genes are ancestrally operating under purifying selection because other forms of selection can be difficult to distinguish (*e.g.,* differentiating relaxed selection from positive selection in asexuals; see Wertheim, et al. 2014).

**Table 2.**
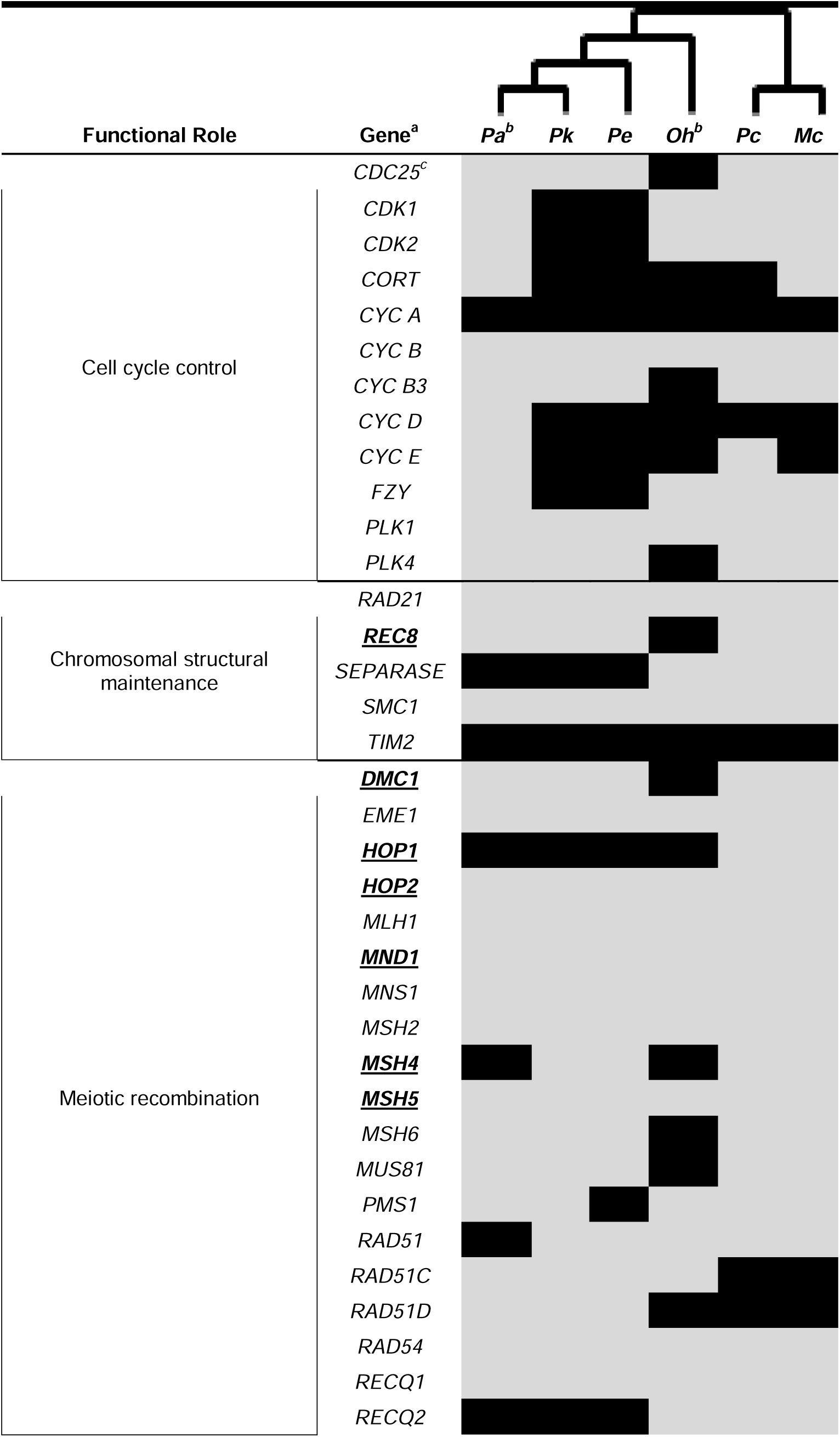
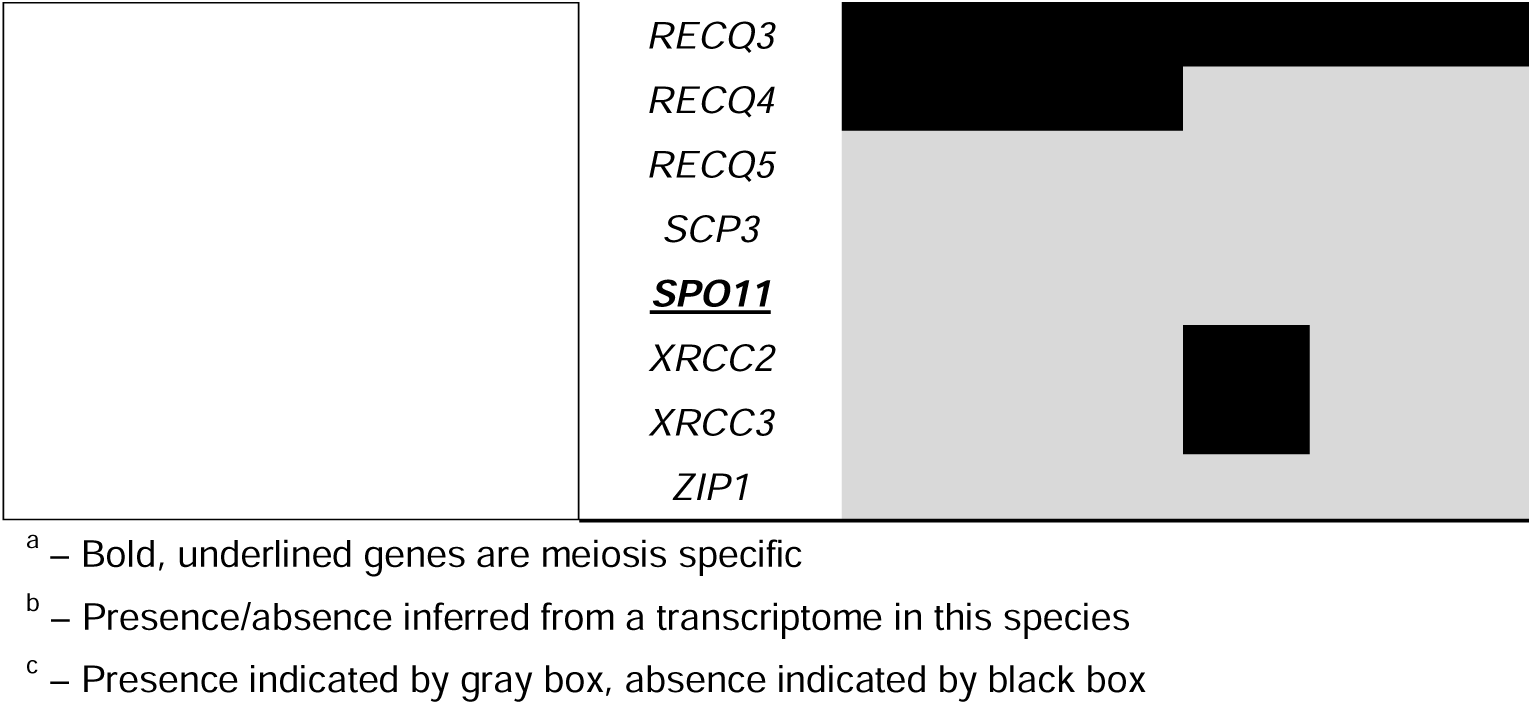
Gene inventory for 44 meiosis genes in six gastropod species.

Here, we establish that the meiosis genes of closely related and sexual *Potamopyrgus* species are highly conserved and nearly identical, establishing the context for which asexual lineages can be compared in the future. We achieved this goal by analyzing the evolution of the 16 meiosis genes that are present in all *Potamopyrgus* and outgroup species. Because the phylogenetic distances between *O*. *hupensis* and the *Potamopyrgus* species are large (see Figure S8), and gene comparisons may be close to or at saturation, we assess both the synonymous substitution rates and protein sequence divergence. We found that all three *Potamopyrgus* taxa are statistically identical concerning substitutions (*p* = 0.993 for synonymous sites; *p* = 0.999 for amino acid distance) when each is compared with the most closely related outgroup, *O. hupensis* (Figure 4). Synonymous distances are possibly saturated for any *O*. *hupensis* to *Potamopyrgus* comparison due to uncorrected medians ranging from 0.701 to 0.712, where a theoretical synonymous distance of 0.75 is functionally indistinguishable from a random alignment of nucleotides. However, saturation does not substantially influencee alignments at the amino acid level, where medians range from 0.152 to 0.155, highlighting the conserved nature of meiosis genes at the protein, but not nucleotide, level. Our findings perfectly align with the expected outcome of meiosis gene evolution in sexual species and now provide a foundation for future studies regarding shifts from purifying to relaxed selection for genes important to sex in *P*. *antipodarum*.

**Figure 4.**
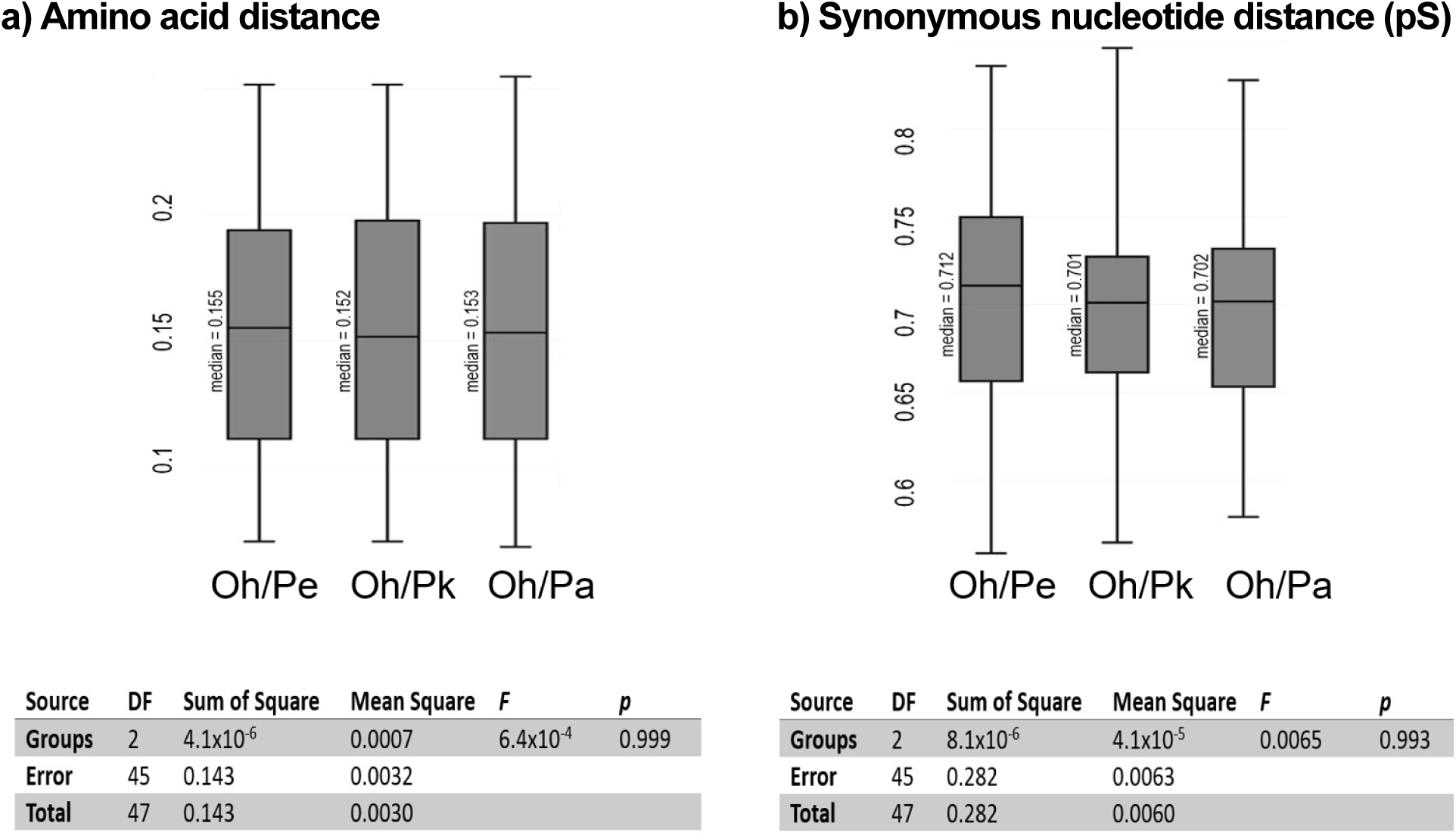
Meiosis gene evolution among *Potamopyrgus* species. (a) Pairwise comparison of amino acid distance from each *Potamopyrgus* species (*P. antipodarum* = Pa, *P. estuarinus* = Pe, *P. kaitunuparaoa* = Pk) to single outgroup *O. hupensis* (Oh) *(p* = 0.999). (b) Pairwise synonymous nucleotide distance (pS) from each *Potamopyrgus* species to *O. hupensis* (*p* = 0.995). Boxplot whiskers extend to minimum and maximum values; box contains Q1 to Q3 and with the median represented by a black line. Statistical significance was assessed using a one-way Anova and Tukey’s range test (α = 0.05). Only meiosis genes with all species represented in single-copy form were used in this analysis.

We also leveraged the hand-curated meiosis-gene alignments as a suitable point of comparison for the genome-wide and automated annotation pipeline we used, MAKER. Automated gene annotation pipelines can produce erroneous annotation, particularly in genomes with complex structures (*e.g.,* polyploidy) and significant repeat content (Bakke, et al. 2009; Ko, et al. 2022; Salzberg 2019). Comparing the carefully hand-annotated meiosis genes to the MAKER versions of those genes can provide an estimate of the reliability and accuracy of our annotations. We find that MAKER transcripts in *P*. *estuarinus* had 21/29 genes that were indistinguishable from their hand-curated counterpart. Five genes had complete and intact reading frames, but MAKER introduced an in-frame sequence (nucleotide insertions that comprised a multiple of three and did not create a premature stop codon) that was not included in the manual annotation. Two genes were split into two separate transcripts, and one gene included non-CDS at either end of the gene sequence while staying in-frame within the gene. The MAKER transcripts in *P*. *kaitunuparaoa* had 14/29 genes that were indistinguishable from their hand-curated counterpart. Eight genes had complete and intact reading frame but also featured instances of both inclusion and deletion of nucleotides with respect to the query. One gene had been split into separate transcripts, and four genes included non-CDS at the ends of the genes. In summary, the MAKER annotations reasonably matched the hand annotations for most meiosis genes, suggesting that the automated approach produced fairly accurate annotations.

### Mitochondrial Genome Assembly and Phylogenetic Analysis

*Potamopyrgus* mitochondrial genomes are comprised of a large single-copy region and a small single-copy region interspersed by a pair of inverted repeats (Sharbrough, et al. 2023). As a result, the non-coding region thought to function as the control region in *Potamopyrgus i*s highly repetitive and intractable with short-read-only approaches. We lacked long reads for *P. kaitunuparaoa* so we were unable to assemble the complete mitochondrial genome. Still, we were able to assemble the large single-copy region for two *P. kaitunuparaoa* and two additional *P. estuarinus* individuals. In these short-read assembles, we found and annotated all 37 genes (13 protein-coding genes, 22 tRNAs, two rRNAs) that are present on the complete and circular reference mitochondrial genome assemblies o*f P. antipodarum* (NC_070577) and *P. estuarinus* (NC_070576), and used this region to evaluate phylogenetic relationships between *Potamopyrgus* species from the mitochondrial perspective, using *O. hupensis* as an outgroup (Figure 5). The mitochondrial genome tree provided strong support for a sister relationship between *P. antipodarum* and *P. kaitunuparaoa*, to the exclusion of *P. estuarinus*.

**Figure 5.**
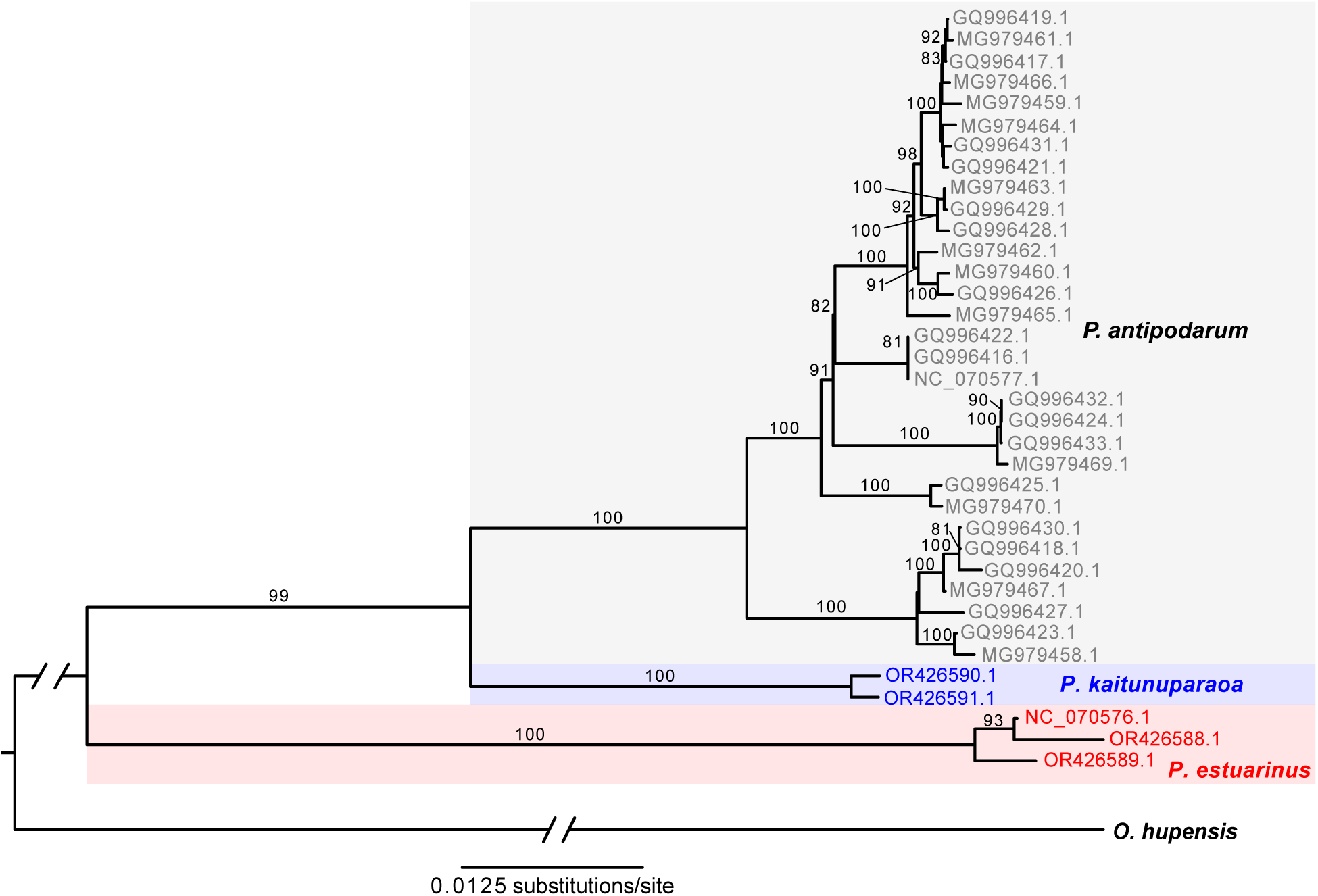
Mitochondrial phylogenetic tree inferred by Maximum Likelihood. *Potamopyrgus kaitunuparaoa* (blue) is sister to *P. antipodarum* (grey) to the exclusion of *P. estuarinus* (red) according to mitochondrial genome data. *Oncomelania hupensis* was used as an outgroup in this analysis.

### Signatures of Introgression Across the Genome

Because phylogenetic analysis using whole mitochondrial genomes revealed that *P. kaitunuparaoa* appears to be sister to *P. antipodarum*, to the exclusion of *P. estuarinus* (the *Pa-Pk* tree) (Figure 5), it was unsurprising that this pattern was also the most common tree topology inferred in gene trees from 1,639 single-copy genes (739 gene trees, or 45.1%). However, a sizable fraction of individual gene trees (478, or 29.2%) instead support a sister relationship between *P. kaitunuparaoa* and *P. estuarinus*, to the exclusion of *P. antipodarum* (the *Pe-Pk* tree). The tree topology featuring *P. estuarinus* and *P. antipodarum* as sister and with *P. kaitunuparaoa* as outgroup (the Pa-Pe tree) was rarer than either of the other two tree topologies (422 gene trees, or 25.7%), likely reflecting baseline levels of incomplete lineage sorting (ILS). The difference in occurrence between the *Pe-Pk* and the *Pa-Pe* trees was even more pronounced when only gene trees with bootstrap support values ≥80 were considered (*Pe-Pk* gene trees = 163, *Pa-Pe* gene trees = 128, binomial test, *p* = 0.0461). We also quantified the three tree topologies in 8,077 1:1:1 triplets that were pruned out of larger orthogroup gene trees to test whether this pattern held in a larger set of genes. Midpoint rooting of triplet gene trees yielded a similar result as above, with 3,326 gene trees supporting the *Pa-Pk* topology (41.2%), 2,658 gene trees supporting the *Pe-Pk* topology (32.9%), and 2,093 gene trees supporting the *Pa-Pe* topology (25.9%). Considering only those gene trees with high bootstrap support (*i.e.,* ≥80), this pattern was even more pronounced, with 1,566 gene trees supporting the *Pa-Pk* topology (45.6%), 1,169 gene trees supporting the *Pe-Pk* topology (34.1%), and only 697 gene trees supporting the *Pa-Pe* topology (20.3%).

The discrepancy in abundance between the two rarer gene tree topologies (*i.e.,* topology in which *P. estuarinus* and *P. kaitunuparaoa* are sister taxa (*Pe-Pk*) is much more common than that in which *P. estuarinus and P. antipodarum* are sister taxa (*Pa-Pe*)) is contrary to the expectations of ILS alone, leading us to hypothesize that some degree of introgression had occurred between these three species. We tested this hypothesis using the commonly employed D-statistic, both in its traditional implementation in a four-taxon arrangement (Durand et al. 2011), as well as in the more recently developed three-taxon method (Hahn and Hibbins 2019). Both the four-taxon method (n = 1,639 1:1:1:1 genes, *D* = 0.1707, 95% CIs: (0.0472 – 0.2849), *Z* = -2.783, *p* = 0.0054) and the three-taxon method (n = 8,077 1:1:1 genes, *D_3_* = 0.0514, 95% CIs: (0.0383 – 0.0629), *Z* = -8.145, *p* < 0.001) identified significant asymmetry across the two least-common tree topologies, indicating that the distribution of gene tree topologies could not be explained by ILS alone (*i.e*., some form of introgression appears to have occurred). The result was similar for all three trimming strategies (*i.e.*, untrimmed, ClipKIT-trimmed, and GBlocks-trimmed) and all three tree topologies (*i.e*., *Pa-Pk*, *Pe-Pk*, and *Pa-Pe*), indicating that some degree of introgression between these taxa appears to have occurred.

To characterize the apparent patterns of introgression, it was first necessary to establish the correct species tree. We reasoned that post-speciation introgression would be expected to result in the following patterns: (1) the introgression tree would be expected to have the shortest patristic distance from outgroup to ingroup compared to the species tree and the ILS tree (species == ILS >> introgression), (2), the introgression trees should be shorter overall than species tree or than the ILS tree if there has been introgression from the ingroup to the outgroup (species == ILS >> introgression), and (3) the branch leading to the midpoint-root-selected outgroup (*i.e*., excluding terminal branches of the ingroup) should be shorter in unrooted introgression trees than in unrooted species or ILS trees (species == ILS >> introgression). In agreement with these predictions, we assigned tree topology for single-copy 1:1:1 triplet trees using midpoint rooting and found that outgroup-to-ingroup patristic distance was significantly lower in *Pe-Pk* trees than in *Pa-Pk* and *Pa-Pe* trees (Figure S9; pairwise Mann-Whitney *U* tests with Holm correction, *p* < 0.0001). Similarly, total tree length was significantly shorter in *Pe-Pk* triplet trees (median = 0.0434 changes per site) than in *Pa-Pk* or *Pa-Pe* trees (pairwise Mann Whitney *U* tests with Holm correction, *p* < 0.0001; Figure S10), and outgroup branch length was shorter in *Pe-Pk* trees than in *Pa-Pk* or *Pa-Pe* trees (pairwise Mann-Whitney *U* tests with Holm correction, *p* < 0.0001; Figure S11). Thus, all three analyses indicate that the mitochondrial tree topology appears to reflect the true species branching order, and that introgression between *P. estuarinus* and *P. kaitunuparaoa* appears to have occurred more recently in the past than the split between the freshwater (*P. kaitunuparaoa* and *P. antipodarum*) and estuarine (*P. estuarinus*) species.

We also tested whether a ghost lineage (*i.e*., an unsampled/extinct *Potamopyrgus* lineage) could be responsible for this signature of introgression. If *Pe-Pk* trees are the result of introgression between *P. antipodarum* and an unsampled ghost lineage rather than introgression between *P. estuarinus* and *P. kaitunuparaoa*, then we would expect a similar distance between *P. estuarinus* and *P. kaitunuparaoa* in both *Pa-Pk* and in *Pe-Pk* gene trees. Instead, we found that the distance between *P. estuarinus* and *P. kaitunuparaoa* is substantially and significantly reduced in *Pe-Pk* trees compared to *Pa-Pk* trees (Mann-Whitney *U* = 3482938, *p* < 0.001, Figure S12), which is inconsistent with a ghost lineage driving the high preponderance of *Pe-Pk* trees. As such, we can conclude that the true species branching order reflects a sister relationship between *P. antipodarum* and *P. kaitunuparaoa*, with some degree of bidirectional introgression occurring between *P. estuarinus* and *P. kaitunuparaoa*.

Although the majority gene tree does appear to be the true species branching order in these species, a sizeable fraction of the *P. estuarinus* and *P. kaitunuparaoa* genomes exhibit an evolutionary history incongruent with that of the mitochondrial genome. Because interactions between the nuclear genome and the mitochondrial genome are likely to contribute to patterns of co-introgression (Beck, et al. 2015) or co-non-introgression (Sharbrough, et al. 2017) of nuclear-encoded genes whose products are targeted to the mitochondria (*i.e*., N-mt genes), we performed enrichment analyses to test whether N-mt genes are enriched for the mitochondrial topology. We used reciprocal best BLASTp hits against the *Drosophila melanogaster* proteome to identify 319 genes in *P. estuarinus* and 293 genes in *P. kaitunuparaoa* that appear to be involved in mitochondrial-nuclear enzyme complexes (*i.e*., OXPHOS complexes I, III, IV, V, the mitoribosome, and mt-aaRS), of which 121 genes were represented in our set of 8,077 1:1:1 triplets (Table S4). Among these 121 genes, 59 had gene tree topologies matching the species tree (*i.e*., *Pe-Pk*), 32 had gene tree topologies that matched the introgression tree (*i.e*., *Pe-Pk*), and 30 genes matched the ILS topology (*i.e*., *Pa-Pe*), a pattern that was not different from the genome-wide pattern (Fisher’s Exact Test *p* = 0.1944; Table 3). When we restricted the analysis to include only the 45 gene trees with ≥80 bootstrap support, we found significantly fewer *Pe-Pk* trees than the genome-wide patterns would predict (Fisher’s Exact Test *p* = 0.0043; Table S5), potentially indicating that mito-nuclear interactions represent a barrier to gene flow between *P. estuarinus* and *P. kaitunuparaoa*. Investigating individual enzyme complexes, only Complex I (*i.e.*, NADH ubiquinone oxidoreductase) exhibited patterns of gene tree topologies that were significantly different from genome-wide patterns, with nine genes (82%, Fisher’s Exact Test *p* = 0.0362) featuring the *Pa-Pk* topology, and only one gene each that exhibited the *Pe-Pk and Pa-Pe* topologies (Table 3). All three Complex I genes with ≥80 bootstrap support exhibited the species topology. Notably, mitochondrially encoded subunits of Complex I exhibit the highest degree of divergence between *P. estuarinus* and *P. kaitunuparaoa* (∼12.5%) of all the mitochondrially encoded gene products (Table S6), potentially indicating that increased mitochondrial genome divergence may have acted as a barrier to interspecific gene flow between *P. estuarinus* and *P. kaitunuparaoa*. This finding echoes recent work in flies (Camus, et al. 2023) and fish (Moran, et al. 2023) that Complex I appears to play an outsized role in hybrid incompatibilities. Because Complex I is the largest single contributor of reactive oxygen species (ROS) to the cell, dysfunction in this complex resulting from cytonuclear incompatibilities may result in elevated and potentially harmful ROS levels in hybrids compared to parental species (Barreto and Burton 2013; Chang, et al. 2016; Du, et al. 2017), providing a potential mechanism for how gene flow could be impeded specifically for nuclear-encoded genes involved in Complex I, but not for other classes of nuclear genes.

**Table 3.** Gene tree topologies in nuclear-encoded mitochondrially targeted genes by enzyme class.

| Enzyme Category | Pa-Pk | Pe-Pk | Pa-Pe | FET *p*-value<sup>a</sup> |
| --- | --- | --- | --- | --- |
| Complex I-NADH ubiquinone oxidoreductase | 9 | 1 | 1 | 0.0362* |
| Complex II-Succinate dehydrogenase<sup>b</sup> | 1 | 1 | 0 | 1 |
| Complex III-Ubiquinol-cytochrome C oxidoreductase | 3 | 1 | 0 | 0.4683 |
| Complex IV-Cytochrome c oxidase | 14 | 12 | 10 | 0.9549 |
| Complex V-F<sub>0</sub>-F<sub>1</sub> ATP synthase | 5 | 3 | 3 | 1 |
| Mitoribosome | 25 | 14 | 15 | 0.5413 |
| aaRS | 2 | 0 | 1 | 0.4881 |
| **Total** | **58** | **31** | **30** | **0.1944** |
| Rooted Triplets | 3326 | 2658 | 2093 | - |

<sup>a</sup> – Fisher's Exact Test was performed against the genome-wide topology counts for rooted triplets, with *p*-values adjusted using the Holm procedure for the Bonferroni correction for multiple comparisons.
<sup>b</sup> – Complex II is entirely nuclear encoded and does not feature any direct interactions between mitochondrially encoded and nuclear encoded genes or gene products. This complex is included here as a negative control and excluded from the Total.
\* – Significant at the *p* < 0.05 level

Finally, we tested whether any gene functional groups were particularly prone to introgression over others, but none of the GO terms were significantly enriched compared to background rates. Additional population-level statistics (*e.g.*, *F_ST_*) will provide a more rigorous test of whether introgression between *P. estuarinus* and *P. kaitunuparaoa* was, at least in part, adaptive.

## Conclusions

The genomes of *P. estuarinus* and *P. kaitunuparaoa* have revealed important insights for understanding the evolution of these biologically interesting species. First, both genomes are similar in size and gene content. This observation provides a clear estimate of the ancestral state of their unique and scientifically prominent congener, *P. antipodarum,* allowing a better understanding of the recent changes in sex, invasiveness, and other derived features in this species. Second, it is clear that *P. kaitunuparaoa* is actually the closest relative to *P. antipodarum*. However, *P. estuarinus* and *P. kaitunuparaoa* have undergone considerable exchange of many nuclear genes more recently than their species divergences from each other and with *P. antipodarum,* but with a strong bias against nuclear-encoded genes with mitochondrial function. The presented study adds to evidence from many systems for the evolutionary importance of genomic incompatibilities between nuclear and mitochondrial genomes.

## Materials and Methods

### Sampling, ploidy estimation, and sequencing

Live specimens of *P. estuarinus* were collected in 2018 from Waikuku Beach estuary in Christchurch, New Zealand, by dragging nets along the surface of the mud in shallow water (<0.5 m depth) and returned alive to the University of Iowa. The *P. kaitunuparaoa* samples were collected in February,2020, from the Mōkau River, New Zealand, where it is sympatric with *P. antipodarum* and *P. estuarinus* (Figure 1).

For *P. estuarinus*, we conducted both DNA and RNA extraction for genome assembly and transcriptome analysis, respectively (McElroy, et al. 2021). To reduce the sequencing of non-target DNA, we isolated snails for several days without feeding them in order to ensure that they had evacuated their guts prior to being dissected for DNA extraction. We then dissected body tissue away from the shell for 22 females, snap-froze this tissue in liquid nitrogen, and 1) shipped the frozen tissue from 20 snails to the Arizona Genomics Institute for DNA extraction and sequencing on the Sequel II (Pacific Biosciences) instrument and 2) extracted genomic DNA for Illumina sequencing from the remaining two snails using a guanidinium thiocyanate and phenol-chloroform-based extraction method (McElroy, et al. 2021; Sharbrough, et al. 2023).

For PacBio sequencing, DNA was extracted by a modified CTAB protocol optimized to deal with common snail contaminants such as mucopolysaccharides (Arseneau, et al. 2017). Extracted high-molecular weight (HMW) DNA was size checked on pulsed-field gel electrophoresis and the sequencing library was constructed following the manufacturer’s protocols using SMRTbell Express Template Prep kit 2.0. The final library was size selected on a BluePippin (Sage Science, Beverly, USA). The recovered library was quantified with Qubit HS and size checked on Femto Pulse (Agilent). Sequencing was performed on Pacbio Sequel II using all standard protocols from the manufacturer.

We also sequenced the *P. estuarinus* and *P. kaitunuparaoa* genomes using Illumina short reads. Briefly, DNA was extracted from individual snails using a CHAOS extraction method (following McElroy, et al. 2021; Sharbrough, et al. 2023). Both sequencing libraries were prepared using a Kapa Hyper Plus preparation, and sequenced with Illumina short-insert 2x150 bp paired-end (PE) reads on an Illumina HiSeq (*P. estuarinus*) and on a NovaSeq 6000 using S4 chemistry (*P. estuarinus* and *P. kaitunuparaoa*) at the Iowa Institute of Human Genetics (IIHG). All sequencing data, assemblies, and annotations have been deposited at NCBI under BioProject PRJNA717745.

### Genome Assembly and Assessment

First, we used our Illumina short-read data to characterize the genomes of *P. estuarinus* and *P. kaitunuparaoa*. Specifically, we estimated genome size using GenomeScope 2.0 (Ranallo-Benavidez, et al. 2020), heterozygosity from k-mer distributions (k=21) generated by Jellyfish (v2.3.0 Marçais and Kingsford 2011), and inferred repeat content using RepeatModeler2 (Flynn, et al. 2020) for both species’ genomes. This information was then used as priors to guide our genome assembly pipeline and parameters.

Genome assembly was conducted using distinct methodologies because the two focal species were sequenced using different technologies, though components of individual inputs did overlap. For *P. estuarinus*, following receipt of raw data in the form of BAM files from the Arizona Genomics Institute, we extracted FASTA and FASTQ files using the bam2fastx utility (https://github.com/PacificBiosciences/bam2fastx), and we used Canu v.1.8 (Koren, et al. 2017) to assemble the first genome draft. Because the sequenced material was derived from 20 individual snail genotypes, default parameters were insufficient to generate either a contiguous or biologically complete assembly. To overcome assembly degradation issues related to large amounts of heterozygosity in the DNA pool, we used an iterative correction of the reads *in silico* to reduce the diversity in raw data. A standard assembly using Canu will use a single iteration of correction, wherein the stochastic errors that arise during sequencing are removed through a consensus decision. We repeated the correction and subsequent assembly steps until we found an asymptote in assembly length, contiguity, and biological completeness as assessed through BUSCO v5.4.2 (Manni, et al. 2021) completeness score. We conducted a total of six correction steps before this asymptote was reached, and adding the seventh iteration did not qualitatively change the results of the three aforementioned metrics.

In the case of *P. kaitunuparaoa*, following receipt of raw data in compressed FASTQ format from IIHG, we used FASTQC (https://www.bioinformatics.babraham.ac.uk/projects/fastqc/) to evaluate read quality and detect adapter contamination. Low-quality reads and adapter contamination were removed using fastp (Chen, et al. 2018). We used a second round of FASTQC to verify read trimming was successful. In order to obtain the most accurate assembly, we used both MaSuRCA and SPAdes assemblers and then merged these independent assemblies using the Purge Haplotig approach (Roach, et al. 2018). In order to improve the contiguity of our assembly, we used the tool RagTag (Alonge, et al. 2022) to conduct a comparative scaffolding using our much more contiguous *P. estuarinus* assembly. Gap filling for both assemblies was done using a combination of Racon (Vaser, et al. 2017) and Pilon (Walker, et al. 2014).

Once both genomes were assembled, we used BUSCO to reevaluate biological completeness in both genomes, compared to the metazoa_odb10 BUSCO reference gene set. To identify contigs derived from contamination, we employed the BlobToolKit v.4.1.4 (Challis, et al. 2020) pipeline with default parameters. This pipeline uses NCBI blastn alignments (Altschul, et al. 1997) to the NCBI nucleotide database and DIAMOND blastx alignments (Buchfink, et al. 2021) to the NCBI non-redundant protein (nr) databases to assign taxonomic affiliations to each contig with an e-value of 10^-25^. BlobToolKit also assessed genome completeness and coverage using results from BUSCO evaluation and BWA-MEM aligned sequencing reads (Li 2013). This process was done for both *P. estuarinus* and *P. kaitunuparaoa* genome assemblies and was visualized in snail- and blob-plots. To correct for contamination, scaffolds identified to be bacterial, viral, or fungal were removed from the assembly.

### Genome Annotation

We used RepeatModeler2 (Flynn, et al. 2020) to build models of repetitive sequence families and masked the assembly using RepeatMasker v.4.0.7 (Smit, et al. 2013) classifying the following transposable element families: “DNA”, “RC”, “LTR”, “LINE”, and “SINE”. We then performed three rounds of MAKER2 v.2.31.9 (Holt and Yandell 2011) annotation to predict genes in the *P. estuarinus* genome assembly. We started with evidence-based input data, followed by two iterative rounds of *ab initio* training based on the previous round(s) of results. As input for the first round of MAKER, we used assembled transcriptomes from three distinct *P. estuarinus* samples (female with embryos, female without embryos, and male) that we combined and clustered with CD-HIT-EST v.4.7 (Fu, et al. 2012; Li and Godzik 2006) to reduce redundancy. Additionally, we used protein models from the *Pomacea canaliculata* (GCF_003073045.1) annotation (Liu, et al. 2018) and translated open reading frames from *Oncomelania hupensis* (SRR1284718) and *P. antipodarum* (SRR6470170) transcriptomes. Next, we trained SNAP v.0.15.4 (Korf 2004) and AUGUSTUS v. 2.5.5 (Stanke, et al. 2006) on the gene models resulting from the transcript and protein-based predictions. To train AUGUSTUS, we ran BUSCO (v.3.0.2; metazoa_odb9 ancestral group) with AUGUSTUS (2.5.5) training enabled and the “--long parameter”. We then reran MAKER, using the SNAP and AUGUSTUS gene models, then repeated the training process and ran a third and final round of MAKER following the same approach as the previous round.

In addition to the RepeatModeler-based results used on the genome assemblies, we compared the repeat content of these genomes based on the reads to account for different assembly qualities. We used dnaPipeTE (v1.3.1c; Goubert, et al. 2015), which assembles consensus repeat sequences with Trinity (v2.5.1; Grabherr, et al. 2011; Haas, et al. 2013) and then classifies the sequences with RepeatMasker (v4.0.7) based on the Repbase libraries (20170127) and estimates repeat family genomic abundance from blastn (BLAST+ v2.2.28) hits of reads against classified repeats. The dnaPipeTE pipeline uses a subsample of reads; here, we sampled reads to 0.20x genome coverage for each species (497000000 for *P. estuarinus* and 510000000 for *P. kaitunuparaoa*). Finally, we used dnaP_utils (https://github.com/clemgoub/dnaPT_utils) to compare the dnaPipeTE results for the two species.

We functionally annotated the completed genome assemblies using DIAMOND Blastx v.2.0.15 (Buchfink, et al. 2021) with the following parameters: e value=1e-5, max hsps=20, very sensitive mode, and searched the predicted transcripts against NCBI’s non-redundant (nr) database. Gene Ontology (GO) terms were then assigned using Blast2GO Basic (v.5.2.5) with default parameters.

### Comparative Genomic Assessment of Assembly and Annotation Quality

To evaluate the quality of our assembly and its annotations, we employed a comparative phylogenetic approach incorporating currently available Caenogastropod genomic resources: *Oncomelania hupensis* (tropical freshwater snail – transcriptome), *Marisa cornuarietis* (common apple snail – genome), and *Pomacea canaliculata* (golden apple snail – genome). These three species are the closest related gastropods to *P. estuarinus* and *P. kaitunuparaoa* for which high-quality genomic/transcriptomic resources are publicly available. Including other gastropods allows us to determine whether the presence or absence of a gene is unique to the *Potamopyrgus* genus or is a common feature of the gastropods of which sequencing is available. Genomes of *M*. *cornuarietis* (GCA_004794655.1) and *Po*. *canaliculata* (GCF_003073045.1) were downloaded from NCBI. To create the *O. hupensis* transcriptome, we downloaded paired-end Illumina RNA-seq reads for *O. hupensis* and *P. antipodarum* from the SRA database (SRR1284718 and SRR6470170, respectively) and assembled them into a transcriptome using Trinity (v2.6.6; Grabherr, et al. 2011; Haas, et al. 2013) with enabled (-- trimmomatic) for read quality control. We then extracted open reading frames with TransDecoder (v5.50; https://github.com/TransDecoder/TransDecoder) and clustered the translated peptide sequences with CD-HIT (v4.7; Fu, et al. 2012; Li and Godzik 2006). We compared biological completeness for each genome and transcriptome using BUSCO v.5.4.2 (Manni, et al. 2021) and the metazoa_odb10 BUSCO reference gene set.

To investigate patterns of evolution within *Potamopyrgus* (and to evaluate the quality of our genomes via comparative genomics), we performed global analyses of phylogenetic patterns across the *P. estuarinus* and *P. kaitunuparaoa* genomes. To accomplish this goal, we used the protein predictions from the *P. estuarinus* and *P. kaitunuparaoa* MAKER genome annotations, as well as predicted proteomes from the *P. antipodarum* transcriptome, the *O. hupensis* transcriptome, *the M. cornuarietis* genome, and the *Po. canaliculata* genome to identify putative orthologs. Following the approach of Sharbrough *et al*. (Sharbrough, et al. 2022), we ran OrthoFinder v2.3.8 (Emms and Kelly 2019) on primary transcripts of all six proteomes. We then aligned CDS sequences from orthologous groups (orthogroups) by translating CDS sequences into amino acids, aligning amino acid sequences with MAFFT v7.480 (Katoh and Standley 2013), and back-translating amino acid sequences into CDS sequences. Alignments were trimmed using three different methods ranging from low to high stringency: untrimmed (low stringency), ClipKIT v1.2.0 (Steenwyk, et al. 2020) (medium stringency), and GBlocks v0.91b (Castresana 2000) (high stringency). Downstream phylogenomic methods were performed at all three stringency levels. Gene trees were inferred from both amino acid and CDS sequence data. Protein-based trees were inferred by RAxML v8.2.12 (Stamatakis 2014), using the -GTRGAMMAIX model of molecular evolution and 100 bootstrap replicates, while CDS trees used gene-specific models of molecular evolution (inferred by jModelTest2 (Darriba, et al. 2012)). The maximum-likelihood tree was inferred using five random tree starts and 100 bootstrap replicates in PhyML v3.3.20211021 (Guindon, et al. 2010).

To evaluate rates and patterns of evolution across orthologs and identify potentially spurious annotations, we also estimated substitution rates under a variety of parameters using the codeml package as part of the PAML v4.9j (Yang 2007). Briefly, we employed both the model 0 (single ω value for the whole tree) and the model 1 (branch-specific ω values) branch models of codeml (i.e., using CDS sequences), for all three possible tree topologies (see Signatures of Introgression section below) for all genes. For all PAML analyses, we also set the RateAncestor parameter to 1, the getSE parameter to 1, the clean_data parameter to 1, the optimation method to 0 (all branches simultaneously), initial kappa = 2, with kappa estimated as part of the model, and initial omega = 0.4.

### Mitochondrial Genome Assembly and Phylogenetic Analysis

The complete *P. estuarinus* mitochondrial genome was assembled using a combination of Illumina HiSeq, Illumina MiSeq, and PacBio reads as described in Sharbrough et al. (2023). *Potamopyrgus* mitochondrial genomes feature a long inverted repeat interspersed by a large dinucleotide repeat (Sharbrough, et al. 2023), so assembling the complete sequence of the *P. kaitunuparaoa* mitochondrial genome was not possible. Instead, we assembled the non-repetitive portion of the mitochondrial genome (*i.e*., the large single-copy region, see Sharbrough et al., (Sharbrough, et al. 2023) for more details) using MEANGS v1.0 (Song, et al. 2021) for two *P. kaitunuparaoa* individuals and two *P. estuarinus* individuals with the -n flag set to 0.1 (10% subsample of reads used) and otherwise default parameters. Mitochondrial genomes were first auto-annotated using MITOS2 (Donath, et al. 2019) and then manually edited.

After assembling mitochondrial genomes of both species, we aligned whole mitochondrial genomes from *P. estuarinus* (NC_070576, OR426588, OR426589)*, P. kaitunuparaoa* (OR426590, OR426591)*, P. antipodarum* (NC_070577, GQ996416-GQ996433, MG979458-MG979467, MG979469-MG979470), and *O. hupensis* (NC_013073), and manually edited the alignment in MEGA11 (Tamura, et al. 2021). We used jModelTest2 v2.1.10 (Darriba, et al. 2012) to choose the best model of molecular evolution using the best-scoring AICc model, and inferred the maximum-likelihood tree using the fast hill-climbing algorithm implemented by RAxML v8.2.12 (Stamatakis 2014) assuming the GTRGAMMMAIX model of molecular evolution, and assessed tree topology with 100 bootstrap replicates. Tree topology was visually inspected to determine relationships among *Potamopyrgus* species in FigTree v.1.4.4 (https://github.com/rambaut/figtree/releases).

### Analysis of Gene Families Associated with Meiotic Function

A catalog of meiosis genes present in the genomes of sexually reproducing *P. estuarinus* and *P. kaitunuparaoa* provides the basis for studying the maintenance/decay of sex-specific genes in the multiple separately derived obligately asexual lineages that characterize *P*. *antipodarum*. To this end, we generated and curated a meiosis gene set using both the original (Schurko and Logsdon Jr 2008; Villeneuve and Hillers 2001) and the most up-to-date (Berdieva, et al. 2021; Tvedte, et al. 2017) inventory of conserved meiosis genes. These inventories discriminate between genes that specifically function in meiosis and genes that are pleiotropic with respect to function outside of meiosis. A combination of NCBI database searches and previously sequenced meiosis genes (Rice 2015) were used to find homologous meiosis genes. We queried the genomes of *P. estuarinus* and *P. kaitunuparaoa* using BLASTx to locate each gene’s scaffold position. We exported the sequence to Sequencher 5.1 (Gene Codes, Ann Arbor, USA) to manually annotate each gene’s start codon, exons, introns, and stop codon. Each annotation was confirmed by 1) comparison to the *P. antipodarum* transcriptome and 2) gene identity by using BLAST. Gene presence was defined as a BLAST query covering >70% of the gene and an -e-value <10^-20^ (as in Tvedte, et al. 2017).

We also used the hand-curated meiosis gene inventory as an opportunity to assess the accuracy of transcript models generated by the automated MAKER pipeline. We compared a subset of the MAKER-generated transcripts to the hand-curated models of meiosis genes. We aligned the annotated meiosis genes derived from *P*. *antipodarum* transcriptomes to compare each of the 29 meiosis genes present in all three focal *Potamopyrgus* species. We generated a local BLAST database for the MAKER transcripts (for both *P*. *estuarinus* and *P*. *kaitunuparaoa*) to make these calls. Then, we used the annotated meiosis genes as queries for BLASTn sequence retrieval. Those retrieved sequences were aligned and scored on the basis of nucleotide identity and whether sequence gaps were introduced in MAKER vs. annotated genes. We scored both nucleotide identity (whether aligned nucleotides match) and gap inclusion (whether nucleotide sequence was introduced or missing) in the MAKER genes. The proportion of genes that exactly or closely match the hand-curated annotations compared with genes that have errors gives insight into the quality of the *P*. *estuarinus* and *P*. *kaitunuparaoa* genome assemblies and the accuracy of our annotation pipeline approach. To understand the phylogenetic context for meiosis gene presence or absence, we also compared meiosis gene toolkits with the transcriptomes and genomes of *O. hupensis*, *M. cornuarietis*, and *Po. canaliculata*.

### Signatures of Introgression Across the Genome

To evaluate phylogenetic relationships between the three *Potamopyrgus* species (*P. antipodarum* (Pa), *P*. *estuarinus* (Pe), and *P*. *kaitunuparaoa* (Pk)), we first extracted 1:1:1 triplets from orthogroup trees using a custom Python script (rootedTriplets.py). Because these triplet relationships were rooted by the other sequences in the alignment, we initially quantified the number of gene trees that supported each of the three possible gene tree topologies: Pa-Pe, Pa-Pk, and Pe-Pk (Figure S1). Tree topology and branch lengths were assessed using custom Python scripts that take advantage of the DendroPy package in Python (Sukumaran and Holder 2010).

The rooted triplets provided evidence of significant asymmetry among the two rarer tree classes (Pe-Pk and Pa-Pe); we reasoned that post-speciation introgression might have played a role in the evolution of these three species. Accordingly, we employed several distinct tests of introgression by taking advantage of amino acid alignments, CDS alignments, and tree branch lengths. These tests are all derived from various implementations of the D-statistic (also known as the ABBA-BABA test), which test for deviations of non-species-tree site patterns from 50-50 in four- (Durand, et al. 2011) and three-taxon sequence alignments (Hahn and Hibbins 2019). Bootstrapping of D-statistics was performed at the gene level to avoid effects of pseudoreplication that result from linkage. Because ghost lineages can contribute to false inferences of introgression between species (Tricou, et al. 2022), we also tested whether a potential ghost lineage could explain our tree topology patterns.

In order to determine whether the species tree reflected the mitochondrial gene tree (*i.e., Pa-Pk* topology) versus the second most-common tree topology (i.e., *Pe-Pk* topology), we developed several predictions for our triplet trees that would enable us to differentiate between various scenarios. First, we expected the introgression tree to exhibit the shortest patristic distance from outgroup to ingroup compared to the species tree and the incomplete-lineage-sorting (ILS) tree (species == ILS >> introgression). Second, the overall length of the triplet trees should be shorter for the introgression trees than for the species or ILS trees (species == ILS >> introgression). After selecting an outgroup using midpoint rooting, the selected outgroup would be expected to have a shorter branch in unrooted introgression trees than in unrooted species or ILS trees (species == ILS >> introgression). We used Mann-Whitney *U* tests to compare these statistics across tree topologies to identify the true species tree. While the mitochondrial genome appeared to follow the inferred species relationship with *P. antipodarum* and *P. kaitunuparaoa* as sister taxa (*Pa-Pk*), much of the nuclear genome (*i.e*., introgressed regions) exhibited a more recent common ancestor between *P. kaitunuparaoa* and *P. estuarinus* (*Pe-Pk*). Because evolutionary mismatches between nuclear-encoded genes that directly interact with mitochondrially encoded genes can produce incompatibilities resulting in reduced fitness, we reasoned that such genes would be particularly resistant to introgression from *P. estuarinus* into *P. kaitunuparaoa*. To test this hypothesis, we reciprocally BLASTed *Drosophila melanogaster* proteins that are involved in mitochondrial-nuclear interactions (*i.e.*, proteins involved in OXPHOS Complexes I, III, IV, and V, the mitoribosome, and mitochondrial tRNA aminoacyl synthetases) against the inferred *P. estuarinus* and *P. kaitunuparaoa* proteomes to identify nuclear-encoded mitochondrially interacting genes in *Potamopyrgus*. We then tested whether tree topologies from these complexes were more likely to match the species topology (*i.e*., *Pa-Pk*) than the rest of the genome using a Fisher’s Exact Test, correcting for multiple comparisons using the Holm procedure for the Bonferroni correction (Holm 1979).

### Data Availability

Raw reads associated with this project are available at NCBI under BioProject PRJNA717745. Assemblies, annotations, and curated gene datasets are available at Zenodo.org (DOI:10.5281/zenodo.10023075). Shell photograph for *P. estuarinus* is available at https://collections.tepapa.govt.nz/object/147506. Shell photograph for *P. kaitunuparaoa* is available at https://collections.tepapa.govt.nz/object/642730. Shell photograph for *P. antipodarum* is available at https://collections.tepapa.govt.nz/object/602110.

## Acknowledgments

We acknowledge funding from USA National Science Foundation grants MCB-1122176 and DEB-1753851, from Iowa Academy of Sciences Grant ISF 17-21 and from Massey University RM22261. JS acknowledges additional funding from Colorado State University and New Mexico Institute for Mining and Technology. We gratefully acknowledge Simon Hills and Mike Winterbourn for help with snail collections and Einat Snir and Keven Knudtson at the Iowa Institute of Human Genetics for assistance with sequencing. We thank J. Bliss and Carson Kephart for snail care. This work utilized the Alpine high performance computing resource at the University of Colorado Boulder. Alpine is jointly funded by the University of Colorado Boulder, the University of Colorado Anschutz, Colorado State University, and the National Science Foundation (award 2201538).

## Notes

### Competing Interest Statement

The authors have declared no competing interest.

