## Supplementary material for "Genome evolution and introgression in the New Zealand mud snails *Potamopyrgus estuarinus* and *Potamopyrgus kaitunuparaoa*": Suppl Materials

**SUPPLEMENTARY FIGURE LEGENDS**

**Figure S1. Possible species tree topologies for *Potamopyrgus* species.** Three possible topologies exist for the relationship between these three species: a sister relationship between *P. antipodarum* and *P. kaitunuparaoa* (Pa-Pk, left), a sister relationship between *P. estuarinus* and *P. kaitunuparaoa* (Pe-Pk, middle), and a sister relationship between *P. antipodarum* and *P. estuarinus* (Pa-Pe, right).

**Figure S2: GenomeScope profile for (a) *P. estuarinus* and (b) *P. kaitunuparaoa*.** Similar genome profiles were obtained for both species from GenomeScope. Both genomes are highly heterozygous (large peak at ~22-27x k-mer coverage). The relatively smaller peak at 50-55x is indicative of homozygous k-mers.

**Figure S3. Genome assembly statistics for *P. estuarinus and P. kaitunuparaoa* genome assemblies.** Snail plots describing genome assembly statistics for *P. estuarinus* (top) and *P. kaitunuparaoa* (bottom) both before (left) and after (right) contaminant removal. The main plot in the center of each panel is divided into 1,000 size-ordered bins, with each bin representing 0.1% of the assembly (total length *P. estuarinus –* 514,633,712 bp; total length *P. kaitunuparaoa –* 596,959,456 bp). Sequence length distribution is shown in dark gray, with the radius scaled to the longest present sequence (red). The arcs in the plot represent the N50 sequence length (orange) and N90 sequence length (pale orange). The pale gray spiral shows the cumulative sequence count on a log scale, with orders of magnitude represented by white scale lines. GC, AT, and N percentages are reflected by the dark blue and pale blue area around the outside of the plot, in the same bins as the inner plot. A summary of BUSCO genes in the metazoa_odb10 dataset are shown in the top right of each plot. Panel A shows the *P. kaitunuparaoa* assembly prior to removal of contaminants, and panel B shows the *P. kaitunuparaoa* assembly following removal of sequences identified with kingdoms in Bamfordvirae, Bacteria-undef, Viridiplantae or Fungi; or phylum in Pseudomonadota or Bacteroidota.

**Figure S4. Blob plots depicting contaminant profiles in *P. estuarinus* and *P. kaitunuparaoa* genome assemblies.** Base coverage plotted as a function of GC proportion in the *P. estuarinus* (top) and (b) *P. kaitunuparaoa* (bottom) genome assemblies before (left) and after (right) contaminant removal. Sequences are coloured by phylum and binned at a resolution of 30 divisions on each axis. Coloured nodes within each bin are sized in proportion to the sum of individual sequence lengths on a logarithmic scale. The assemblies have been filtered to exclude sequences with kingdom in Bamfordvirae, Bacteria-undef, Viridiplantae or Fungi; or phylum in Pseudomonadota or Bacteroidota. Histograms show the distribution of sequence length sums along each axis.

**Figure S5. Distribution of BLAST hits of *P. estaurinus* and *P.* kaitunuparaoa transcripts before and after removal of potential contaminants.** Species distribution of BLAST hits before (a-d) and after (e-h) contaminant removal for *P. estuarinus* (left) and *P. kaitunuparaoa* (right) genome assemblies, obtained using BLAST2GO of assembly transcript annotations. Blue histograms (a, c, e, g) represent all BLAST hits, and green histograms (b, d, f, h) only include the top BLAST hits.

**Figure S6. Homologous gene groups in assemblies from six caenogastropodan taxa.** Upset plot depicting the number of genes fitting into homologous gene groups (group membership specified in bottom panel) for multi-copy genes (top panel) and single-copy genes (bottom panel). Singleton genes from *P. antipodarum* and *O. hupensis* (transcriptomes) that were unique to those transcriptomes are not shown here.

**Figure S7. Estimates of *d_N_ /d_S_* for single-copy orthologs in *P. estuarinus* and *P. kaitunuparaoa*.** Density plot depicting the distribution of *d_N_ /d_S_* estimated from single-copy orthologous gene groups using model 0 of codeml, assuming the *Pa-Pk* tree topology. Outliers are not included in this plot for display purposes.

**Figure S8. Branch-specific estimates of *d_S_* in single-copy orthologs in *P.estuarinus* and *P. kaitunuparaoa*.** Density plots depicting pairwise branch-specific *d_S_* distributions estimated from single-copy orthologs using model 1 in codeml. Distributions are mapped onto the *Potamopyrgus* phylogeny. Branch lengths on the phylogeny reflect branch-specific *d_S_* estimates from concatenated alignment of all single-copy genes. Red lines reflect *d_S_* cutoff points used to filter bad alignments from molecular evolution and phylogenetic analyses.

**Figure S9. Patristic distance of outgroup to ingroup taxa in 1:1:1 triplet gene trees across midpoint-root-determined tree topologies**. Outgroup-to-ingroup patristic distance for *Pa-Pk* trees (grey), *Pe-Pk* trees (blue), and *Pa-Pe* trees (orange) inferred from 1:1:1 triplet gene trees. Statistical groupings (red letters) were determined using a series of pairwise Mann-Whitney U tests, with significance corrected for multiple comparisons using the Holm procedure for the Bonferroni correction.

**Figure S10. Total tree length in 1:1:1 triplet gene trees across midpoint-root-determined tree topologies**. Total tree length for *Pa-Pk* trees (grey), *Pe-Pk* trees (blue), and *Pa-Pe* trees (orange) determined from 1:1:1 triplet gene trees. Statistical groupings (red letters) were determined using a series of pairwise Mann-Whitney U tests, with significance corrected for multiple comparisons using the Holm procedure for the Bonferroni correction.

**Figure S11. Outgroup branch length in 1:1:1 triplet gene trees across midpoint-root-determined tree topologies**. Outgroup branch length for *Pa-Pk* trees (grey), *Pe-Pk* trees (blue), and *Pa-Pe* trees (orange) determined from 1:1:1 triplet gene trees. Statistical groupings (red letters) were determined using a series of pairwise Mann-Whitney U tests, with significance corrected for multiple comparisons using the Holm procedure for the Bonferroni correction.

**Figure S12. Summed branch lengths between *P. estuarinus* and *P. kaitunuparaoa* in 1:1:1 triplet gene trees across midpoint-root-determined tree topologies**. Summed branch lengths from *P. estuarinus* to *P. kaitunuparaoa in Pa-Pk* trees (grey), *Pe-Pk* trees (blue), and *Pa-Pe* trees (orange) in 1:1:1 triplet gene trees. Statistical groupings (red letters) were determined using a series of pairwise Mann-Whitney U tests, with significance corrected for multiple comparisons using the Holm procedure for the Bonferroni correction.
