## Supplementary material for "Genome evolution and introgression in the New Zealand mud snails *Potamopyrgus estuarinus* and *Potamopyrgus kaitunuparaoa*": Suppl Figures

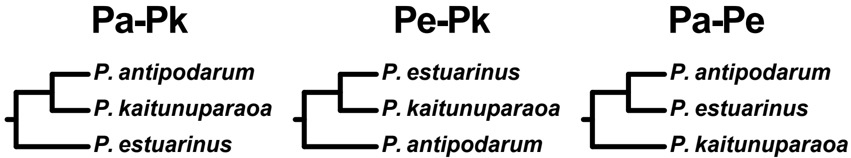
Pa-Pk
P. antipodarum
P. kaitunuparaoa
P. estuarinus
Pe-Pk
P. estuarinus
P. kaitunuparaoa
P. antipodarum
Pa-Pe
P. antipodarum
P. estuarinus
P. kaitunuparaoa

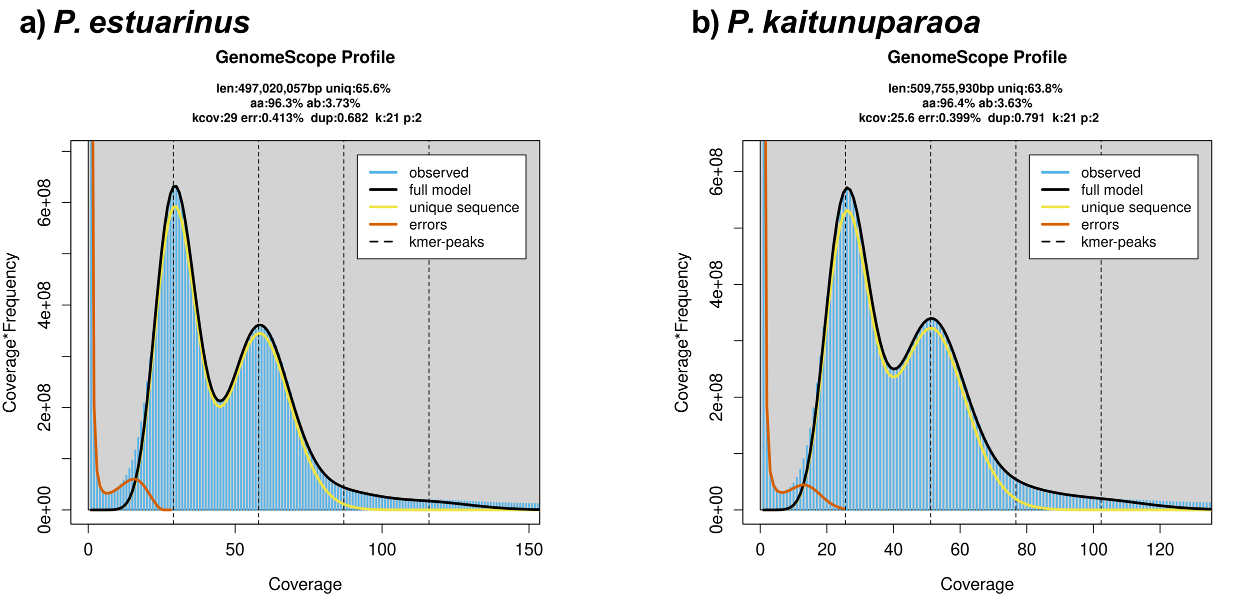

a) P. estuarinus
GenomeScope Profile
len:497,020,057bp uniq:65.6%
aa:96.3% ab:3.73%
kcov:29 err:0.413% dup:0.682 k:21 p:2
observed
full model
unique sequence
errors
kmer-peaks
Coverage*Frequency
0e+00
2e+08
4e+08
6e+08
0
50
100
150
Coverage
b) P. kaitunuparaoa
GenomeScope Profile
len:509,755,930bp uniq:63.8%
aa:96.4% ab:3.63%
kcov:25.6 err:0.399% dup:0.791 k:21 p:2
observed
full model
unique sequence
errors
kmer-peaks
Coverage*Frequency
0e+00
2e+08
4e+08
6e+08
0
20
40
60
80
100
120
Coverage

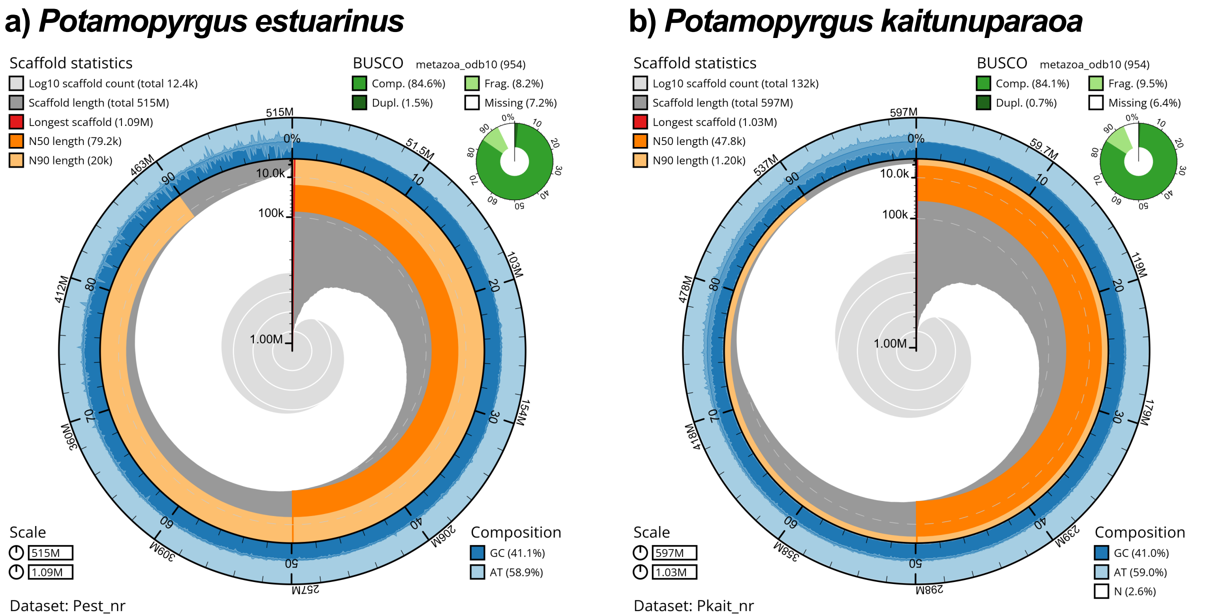

a) Potamopyrgus estuarinus
Scaffold statistics
Log10 scaffold count (total 12.4k)
Scaffold length (total 515M)
Longest scaffold (1.09M)
N50 length (79.2k)
N90 length (20k)
BUSCO metazoa_odb10 (954)
Comp. (84.6%)
Frag. (8.2%)
Dupl. (1.5%)
Missing (7.2%)
Scale
515M
1.09M
Composition
GC (41.1%)
AT (58.9%)
Dataset: Pest_nr
b) Potamopyrgus kaitunuparaoa
Scaffold statistics
Log10 scaffold count (total 132k)
Scaffold length (total 597M)
Longest scaffold (1.03M)
N50 length (47.8k)
N90 length (1.20k)
BUSCO metazoa_odb10 (954)
Comp. (84.1%)
Frag. (9.5%)
Dupl. (0.7%)
Missing (6.4%)
Scale
597M
1.03M
Composition
GC (41.0%)
AT (59.0%)
N (2.6%)
Dataset: Pkait_nr

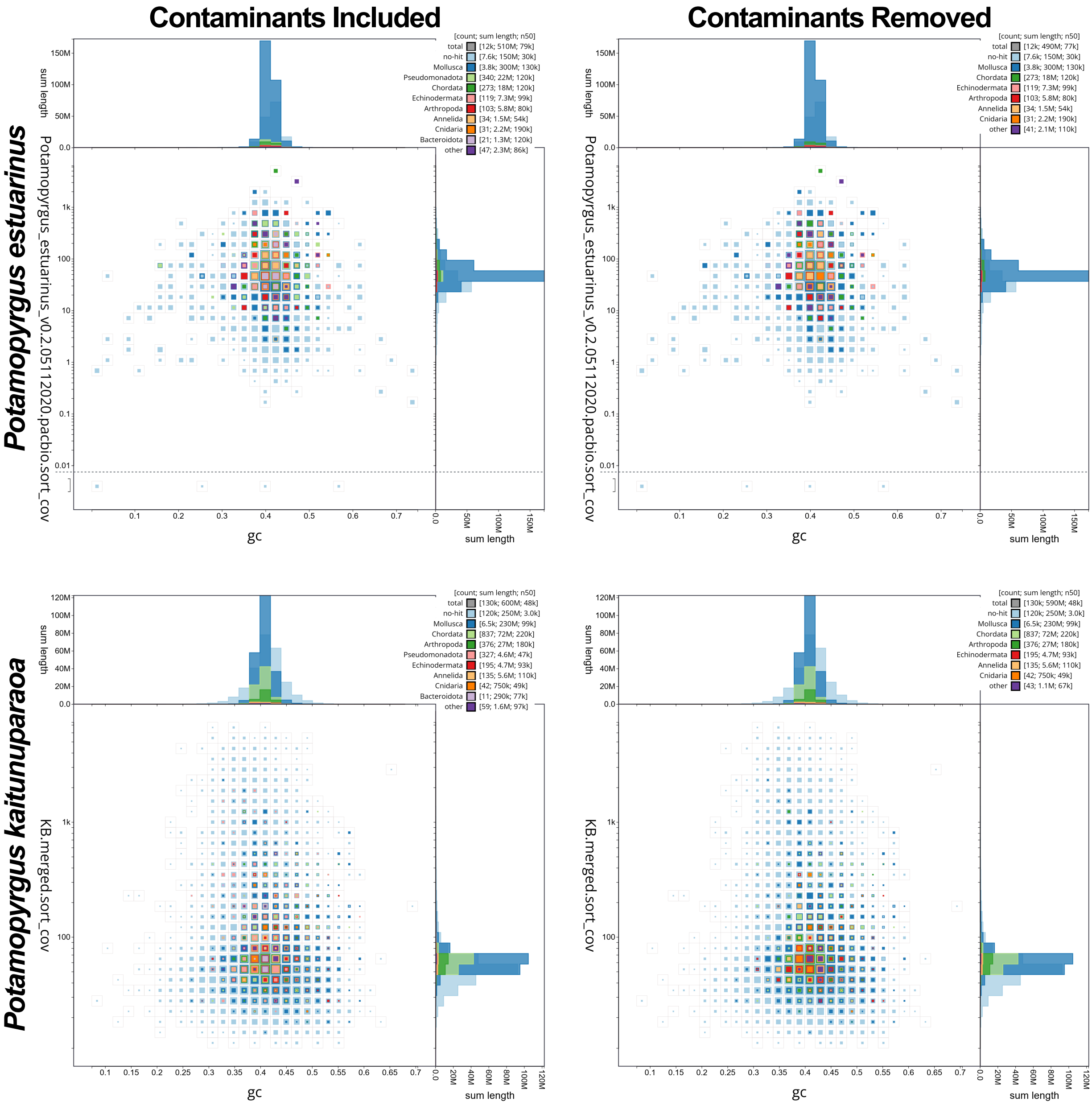

Contaminants Included
Contaminants Removed
Potamopyrgus estuarinus
Potamopyrgus kaitunuparaoa
Potamopyrgus_estuarinus_v0.2.05112020.pacbio.sort_cov
KB.merged.sort_cov
sum length
gc
[count; sum length; n50]
total [12k; 510M; 79k]
no-hit [7.6k; 150M; 30k]
Mollusca [3.8k; 300M; 130k]
Pseudomonadota [340; 22M; 120k]
Chordata [273; 18M; 120k]
Echinodermata [119; 7.3M; 99k]
Arthropoda [103; 5.8M; 80k]
Annelida [34; 1.5M; 54k]
Cnidaria [31; 2.2M; 190k]
Bacteroidota [21; 1.3M; 120k]
other [47; 2.3M; 86k]
total [12k; 490M; 77k]
no-hit [7.6k; 150M; 30k]
Mollusca [3.8k; 300M; 130k]
Chordata [273; 18M; 120k]
Echinodermata [119; 7.3M; 99k]
Arthropoda [103; 5.8M; 80k]
Annelida [34; 1.5M; 54k]
Cnidaria [31; 2.2M; 190k]
other [41; 2.1M; 110k]
total [130k; 600M; 48k]
no-hit [120k; 250M; 3.0k]
Mollusca [6.5k; 230M; 99k]
Chordata [837; 72M; 220k]
Arthropoda [376; 27M; 180k]
Pseudomonadota [327; 4.6M; 47k]
Echinodermata [195; 4.7M; 93k]
Annelida [135; 5.6M; 110k]
Cnidaria [42; 750k; 49k]
Bacteroidota [11; 290k; 77k]
other [59; 1.6M; 97k]
total [130k; 590M; 48k]
no-hit [120k; 250M; 3.0k]
Mollusca [6.5k; 230M; 99k]
Chordata [837; 72M; 220k]
Arthropoda [376; 27M; 180k]
Echinodermata [195; 4.7M; 93k]
Annelida [135; 5.6M; 110k]
Cnidaria [42; 750k; 49k]
other [43; 1.1M; 67k]

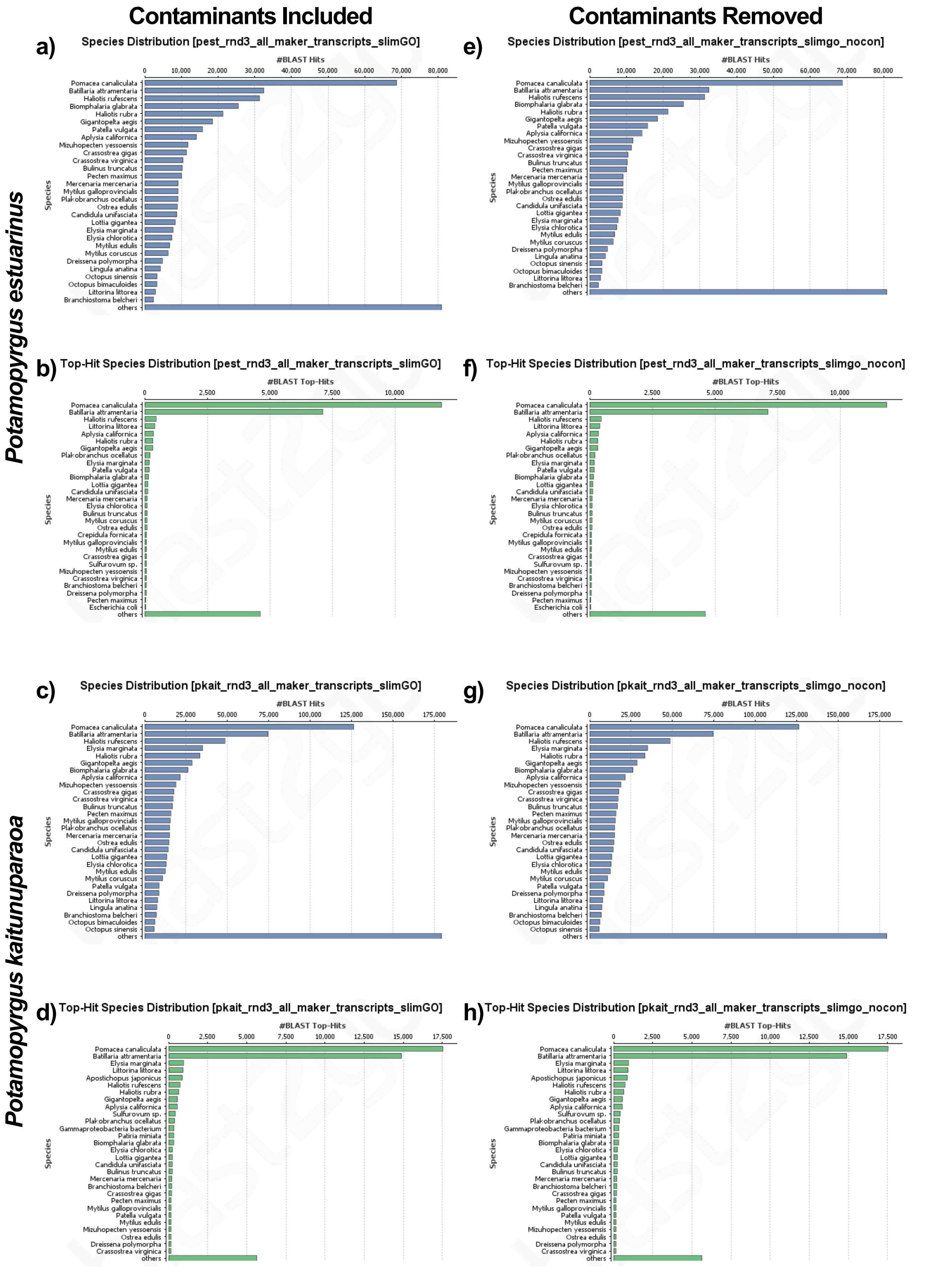

Contaminants Included
Contaminants Removed
Potamopyrgus estuarinus
Potamopyrgus kaitunuparaoa
a)
Species Distribution [pest_rnd3_all_maker_transcripts_slimGO]
#BLAST Hits
0
10,000
20,000
30,000
40,000
50,000
60,000
70,000
80,000
Species
Pomacea canaliculata
Batillaria attramentaria
Haliotis rufescens
Biomphalaria glabrata
Haliotis rubra
Gigantopelta aegis
Patella vulgata
Aplysia californica
Mizuhopecten yessoensis
Crassostrea gigas
Crassostrea virginica
Bulinus truncatus
Pecten maximus
Mercenaria mercenaria
Mytilus galloprovincialis
Plakobranchus ocellatus
Ostrea edulis
Candidula unifasciata
Lottia gigantea
Elysia marginata
Elysia chlorotica
Mytilus edulis
Mytilus coruscus
Dreissena polymorpha
Lingula anatina
Octopus sinensis
Octopus bimaculoides
Littorina littorea
Branchiostoma belcheri
others
e)
Species Distribution [pest_rnd3_all_maker_transcripts_slimgo_nocon]
#BLAST Hits
0
10,000
20,000
30,000
40,000
50,000
60,000
70,000
80,000
Species
Pomacea canaliculata
Batillaria attramentaria
Haliotis rufescens
Biomphalaria glabrata
Haliotis rubra
Gigantopelta aegis
Patella vulgata
Aplysia californica
Mizuhopecten yessoensis
Crassostrea gigas
Crassostrea virginica
Bulinus truncatus
Pecten maximus
Mercenaria mercenaria
Mytilus galloprovincialis
Plakobranchus ocellatus
Ostrea edulis
Candidula unifasciata
Lottia gigantea
Elysia marginata
Elysia chlorotica
Mytilus edulis
Mytilus coruscus
Dreissena polymorpha
Lingula anatina
Octopus sinensis
Octopus bimaculoides
Littorina littorea
Branchiostoma belcheri
others
b)
Top-Hit Species Distribution [pest_rnd3_all_maker_transcripts_slimGO]
#BLAST Top-Hits
0
2,500
5,000
7,500
10,000
Species
Pomacea canaliculata
Batillaria attramentaria
Haliotis rufescens
Littorina littorea
Aplysia californica
Haliotis rubra
Gigantopelta aegis
Plakobranchus ocellatus
Elysia marginata
Patella vulgata
Biomphalaria glabrata
Lottia gigantea
Candidula unifasciata
Mercenaria mercenaria
Elysia chlorotica
Bulinus truncatus
Mytilus coruscus
Ostrea edulis
Crepidula fornicata
Mytilus galloprovincialis
Mytilus edulis
Crassostrea gigas
Sulfurovum sp.
Mizuhopecten yessoensis
Crassostrea virginica
Branchiostoma belcheri
Dreissena polymorpha
Pecten maximus
Escherichia coli
others
f)
Top-Hit Species Distribution [pest_rnd3_all_maker_transcripts_slimgo_nocon]
#BLAST Top-Hits
0
2,500
5,000
7,500
10,000
Species
Pomacea canaliculata
Batillaria attramentaria
Haliotis rufescens
Littorina littorea
Aplysia californica
Haliotis rubra
Gigantopelta aegis
Plakobranchus ocellatus
Elysia marginata
Patella vulgata
Biomphalaria glabrata
Lottia gigantea
Candidula unifasciata
Mercenaria mercenaria
Elysia chlorotica
Bulinus truncatus
Mytilus coruscus
Ostrea edulis
Crepidula fornicata
Mytilus galloprovincialis
Mytilus edulis
Crassostrea gigas
Sulfurovum sp.
Mizuhopecten yessoensis
Crassostrea virginica
Branchiostoma belcheri
Dreissena polymorpha
Pecten maximus
Escherichia coli
others
c)
Species Distribution [pkait_rnd3_all_maker_transcripts_slimGO]
#BLAST Hits
0
25,000
50,000
75,000
100,000
125,000
150,000
175,000
Species
Pomacea canaliculata
Batillaria attramentaria
Haliotis rufescens
Elysia marginata
Haliotis rubra
Gigantopelta aegis
Biomphalaria glabrata
Aplysia californica
Mizuhopecten yessoensis
Crassostrea gigas
Crassostrea virginica
Bulinus truncatus
Pecten maximus
Mytilus galloprovincialis
Plakobranchus ocellatus
Mercenaria mercenaria
Ostrea edulis
Candidula unifasciata
Lottia gigantea
Elysia chlorotica
Mytilus edulis
Mytilus coruscus
Patella vulgata
Dreissena polymorpha
Littorina littorea
Lingula anatina
Branchiostoma belcheri
Octopus bimaculoides
Octopus sinensis
others
g)
Species Distribution [pkait_rnd3_all_maker_transcripts_slimgo_nocon]
#BLAST Hits
0
25,000
50,000
75,000
100,000
125,000
150,000
175,000
Species
Pomacea canaliculata
Batillaria attramentaria
Haliotis rufescens
Elysia marginata
Haliotis rubra
Gigantopelta aegis
Biomphalaria glabrata
Aplysia californica
Mizuhopecten yessoensis
Crassostrea gigas
Crassostrea virginica
Bulinus truncatus
Pecten maximus
Mytilus galloprovincialis
Plakobranchus ocellatus
Mercenaria mercenaria
Ostrea edulis
Candidula unifasciata
Lottia gigantea
Elysia chlorotica
Mytilus edulis
Mytilus coruscus
Patella vulgata
Dreissena polymorpha
Littorina littorea
Lingula anatina
Branchiostoma belcheri
Octopus bimaculoides
Octopus sinensis
others
d)
Top-Hit Species Distribution [pkait_rnd3_all_maker_transcripts_slimGO]
#BLAST Top-Hits
0
2,500
5,000
7,500
10,000
12,500
15,000
17,500
Species
Pomacea canaliculata
Batillaria attramentaria
Elysia marginata
Littorina littorea
Apostichopus japonicus
Haliotis rufescens
Haliotis rubra
Gigantopelta aegis
Aplysia californica
Sulfurovum sp.
Plakobranchus ocellatus
Gammaproteobacteria bacterium
Patiria miniata
Biomphalaria glabrata
Elysia chlorotica
Lottia gigantea
Candidula unifasciata
Bulinus truncatus
Mercenaria mercenaria
Branchiostoma belcheri
Crassostrea gigas
Pecten maximus
Mytilus galloprovincialis
Patella vulgata
Mytilus edulis
Mizuhopecten yessoensis
Ostrea edulis
Dreissena polymorpha
Crassostrea virginica
others
h)
Top-Hit Species Distribution [pkait_rnd3_all_maker_transcripts_slimgo_nocon]
#BLAST Top-Hits
0
2,500
5,000
7,500
10,000
12,500
15,000
17,500
Species
Pomacea canaliculata
Batillaria attramentaria
Elysia marginata
Littorina littorea
Apostichopus japonicus
Haliotis rufescens
Haliotis rubra
Gigantopelta aegis
Aplysia californica
Sulfurovum sp.
Plakobranchus ocellatus
Gammaproteobacteria bacterium
Patiria miniata
Biomphalaria glabrata
Elysia chlorotica
Lottia gigantea
Candidula unifasciata
Bulinus truncatus
Mercenaria mercenaria
Branchiostoma belcheri
Crassostrea gigas
Pecten maximus
Mytilus galloprovincialis
Patella vulgata
Mytilus edulis
Mizuhopecten yessoensis
Ostrea edulis
Dreissena polymorpha
Crassostrea virginica
others

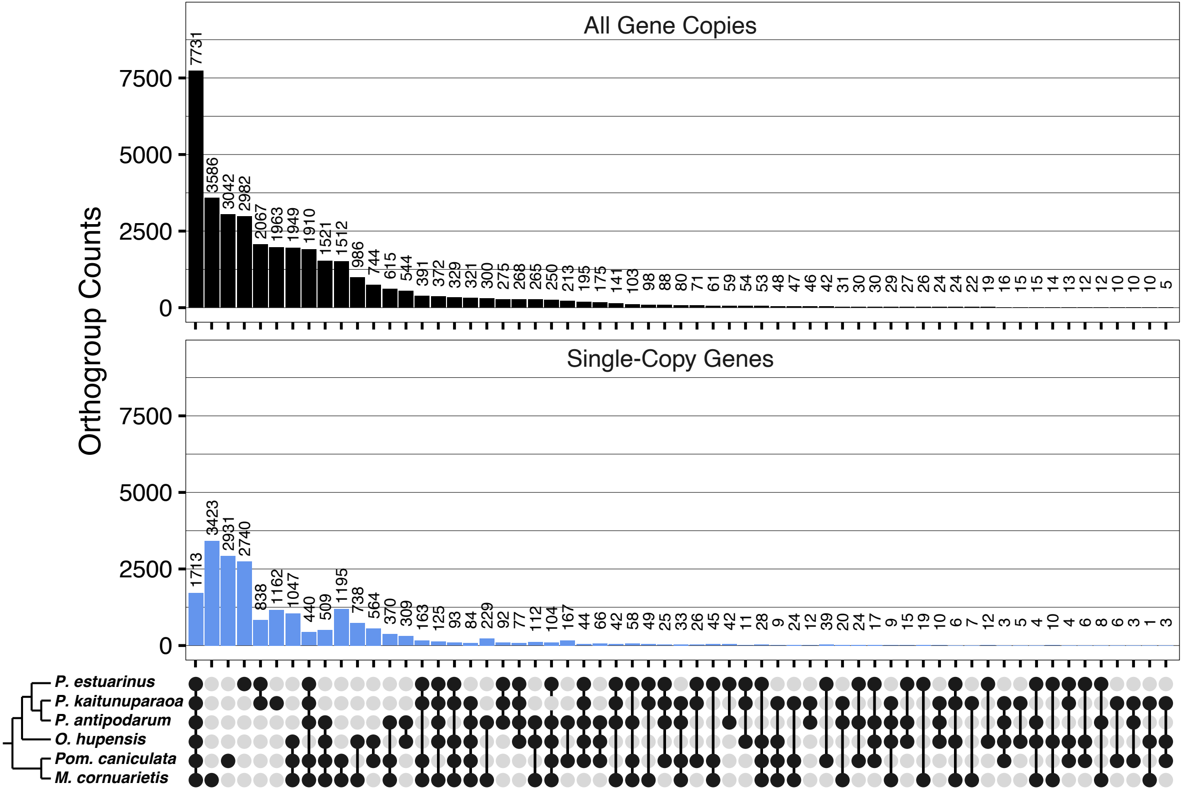
All Gene Copies
Single-Copy Genes
Orthogroup Counts
0
2500
5000
7500
7731
3586
3042
2982
2067
1963
1949
1910
1521
1512
986
744
615
544
391
372
329
321
300
275
268
265
250
213
195
175
141
103
98
88
80
71
61
59
54
53
48
47
46
42
31
30
30
29
27
26
24
24
22
19
16
15
15
14
13
12
12
10
10
10
5
1713
3423
2931
2740
838
1162
1047
440
509
1195
738
564
370
309
163
125
93
84
229
92
77
112
104
167
44
66
42
58
49
25
33
26
45
42
11
28
9
24
12
39
20
24
17
9
15
19
10
6
7
12
3
5
4
10
4
6
8
6
3
1
3
P. estuarinus
P. kaitunuparaoa
P. antipodarum
O. hupensis
Pom. caniculata
M. cornuarietis

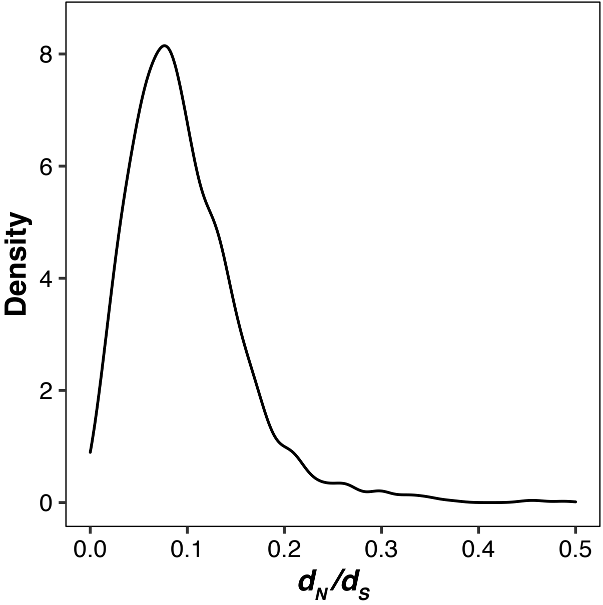

Density
8
6
4
2
0
0.0
0.1
0.2
0.3
0.4
0.5
$d_N/d_S$

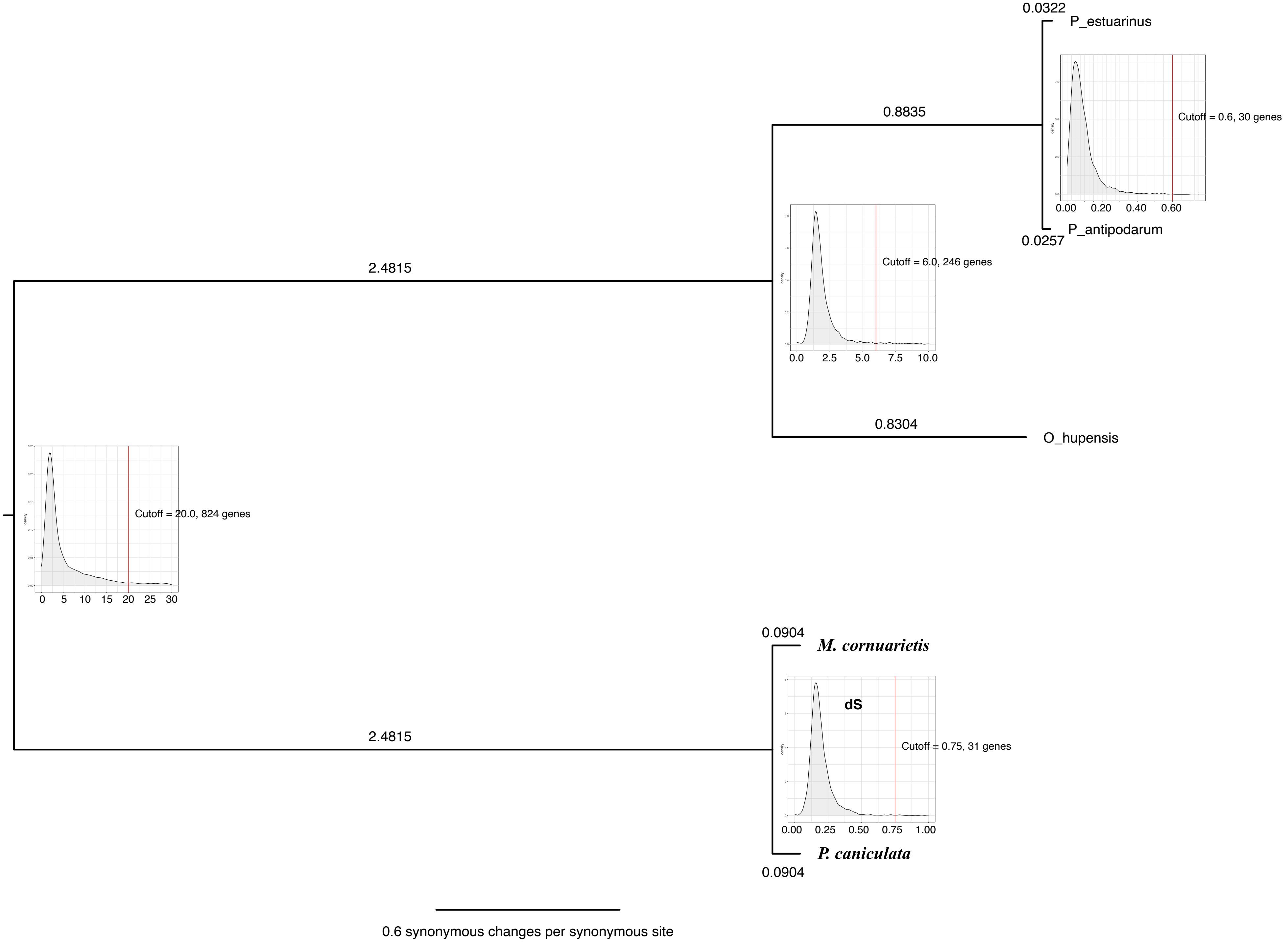

0.0322
P_estuarinus
0.8835
Cutoff = 0.6, 30 genes
0.00 0.20 0.40 0.60
P_antipodarum
0.0257
2.4815
Cutoff = 6.0, 246 genes
0.0 2.5 5.0 7.5 10.0
0.8304
O_hupensis
Cutoff = 20.0, 824 genes
0 5 10 15 20 25 30
0.0904
M. cornuarietis
dS
Cutoff = 0.75, 31 genes
2.4815
0.00 0.25 0.50 0.75 1.00
P. caniculata
0.0904
0.6 synonymous changes per synonymous site

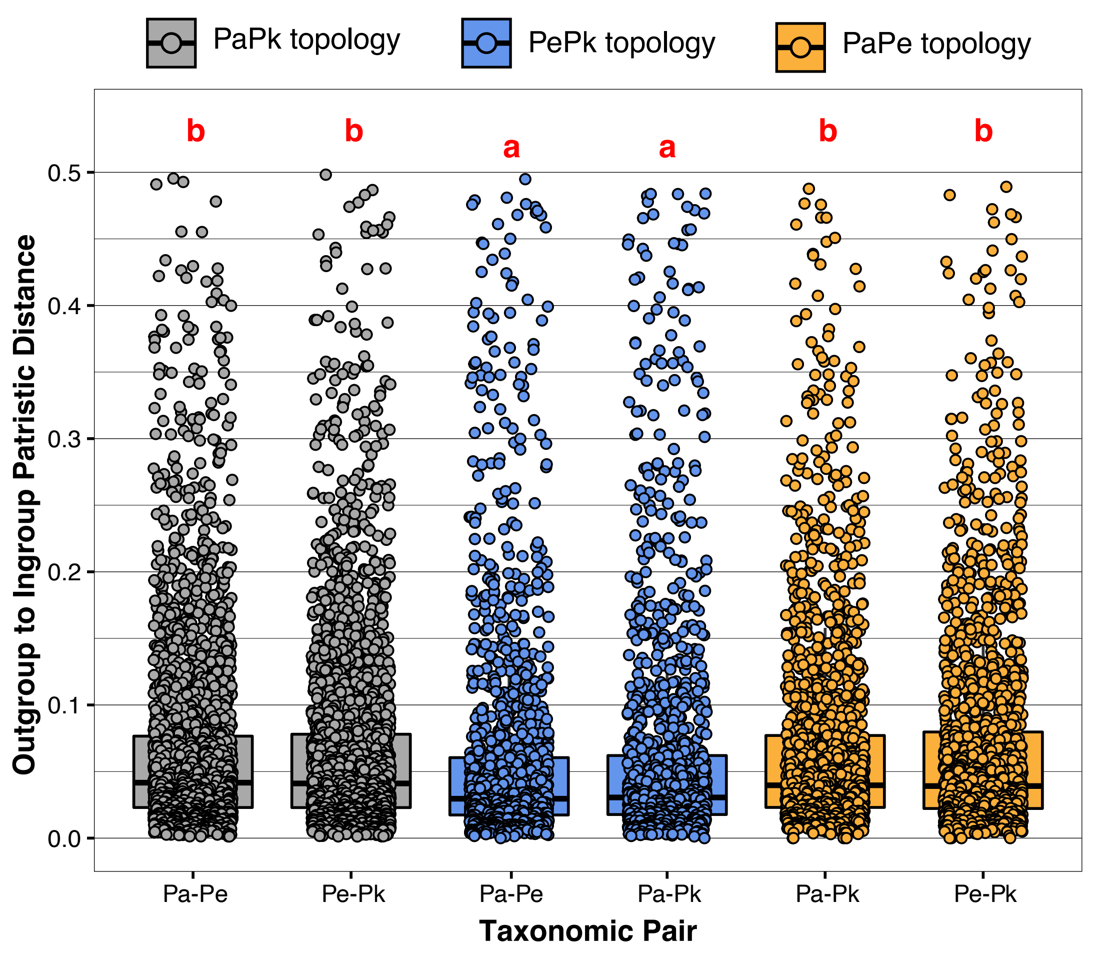
PaPk topology
PePk topology
PaPe topology
b
b
a
a
b
b
0.5
0.4
0.3
0.2
0.1
0.0
Outgroup to Ingroup Patristic Distance
Pa-Pe
Pe-Pk
Pa-Pe
Pa-Pk
Pa-Pk
Pe-Pk
Taxonomic Pair

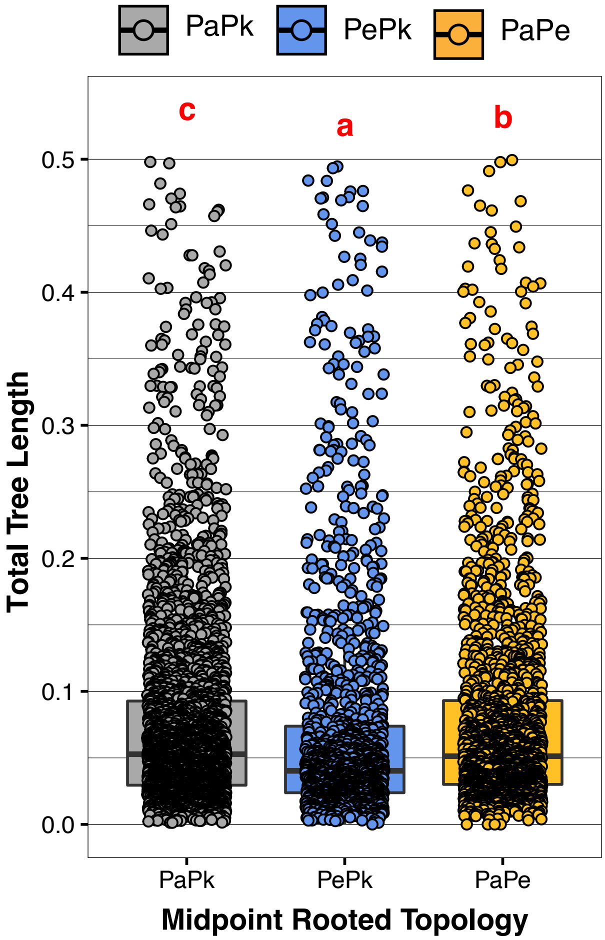
PaPk
PePk
PaPe
c
a
b
Total Tree Length
0.0
0.1
0.2
0.3
0.4
0.5
PaPk
PePk
PaPe
Midpoint Rooted Topology

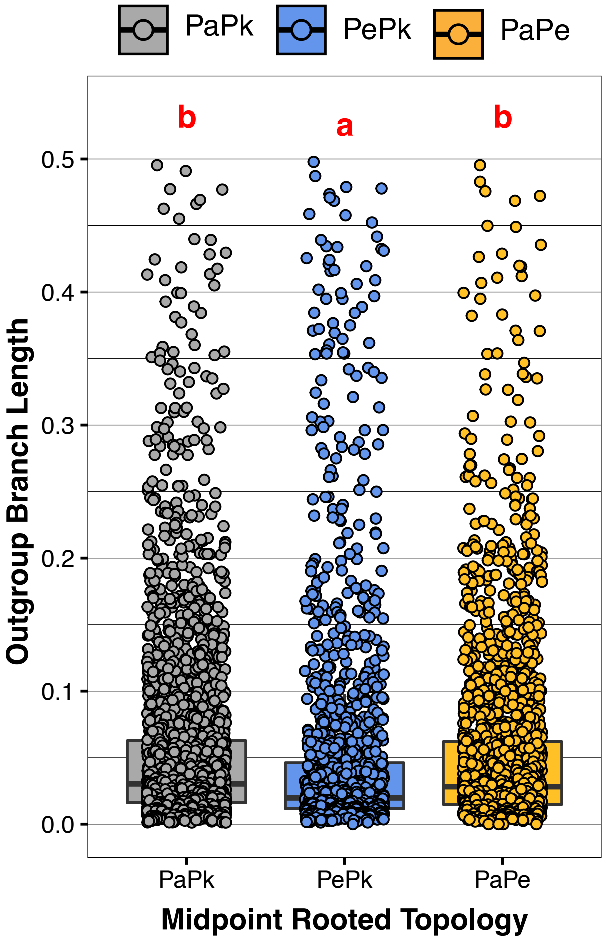

PaPk
PePk
PaPe
b
a
b
0.5
0.4
0.3
0.2
0.1
0.0
Outgroup Branch Length
PaPk
PePk
PaPe
Midpoint Rooted Topology

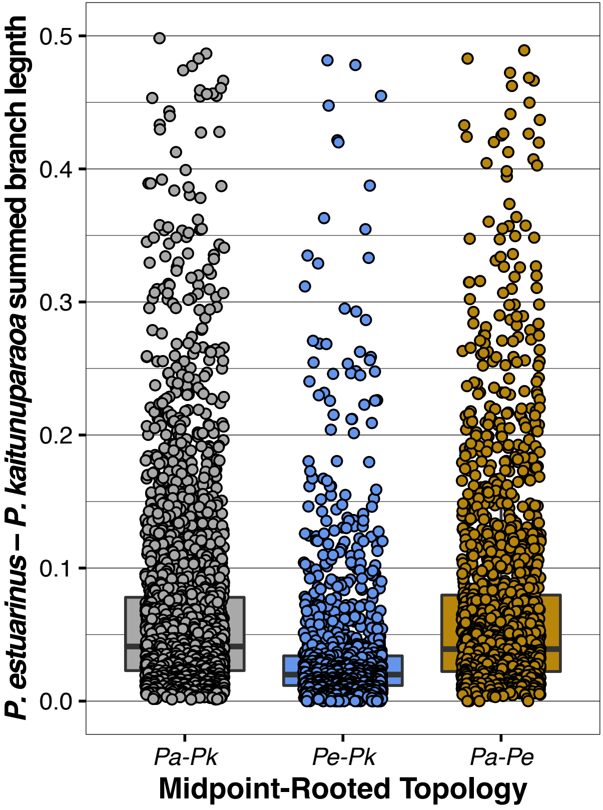

P. estuarinus – P. kaitunuparaoa summed branch legnth
0.0
0.1
0.2
0.3
0.4
0.5
Pa-Pk
Pe-Pk
Pa-Pe
Midpoint-Rooted Topology
